# Placental structure, function and mitochondrial phenotype relate to fetal size and sex in mice

**DOI:** 10.1101/2021.07.22.453249

**Authors:** Esteban Salazar-Petres, Daniela Pereira Carvalho, Jorge Lopez-Tello, Amanda Nancy Sferruzzi-Perri

**Affiliations:** Department of Physiology, Development and Neuroscience, Centre for Trophoblast Research, University of Cambridge, Cambridge, United Kingdom

**Keywords:** Placenta, fetus, sex, mitochondria, transport, hormones

## Abstract

Fetal growth depends on placental function, which requires energy from mitochondria. Here we investigated whether mitochondrial function in the placenta relates to growth of the lightest and heaviest fetuses of each sex within the litter of mice. Placentas from the lightest and heaviest fetuses were taken to evaluate placenta morphology (stereology), mitochondrial energetics (high-resolution respirometry), and mitochondrial regulators, nutrient transporters, hormone handling and signalling pathways (qPCR and western blotting). We found that mitochondrial complex I and II oxygen consumption rate was greater for placentas supporting the lightest female fetuses, although placental complex I abundance of the lightest females and complexes III and V of the lightest males were decreased compared to their heaviest counterparts. Expression of mitochondrial biogenesis (*Nrf1*) and fission (*Drp1* and *Fis1*) genes was lower in the placenta from the lightest females, whilst biogenesis-related gene *Tfam* was greater in the placenta of the lightest male fetuses. Additionally, placental morphology and steroidogenic gene (*Cyp17a1* and *Cyp11a1*) expression was aberrant for the lightest females, but glucose transporter (*Glut1*) expression was lower in only the lightest males versus their heaviest counterparts. Differences in intra-litter placental phenotype were related to sex-dependent changes in the expression of hormone responsive (androgen receptor) and metabolic signalling (AMPK, AKT, PPARγ) pathways. Thus, in normal mouse pregnancy, placental structure, function and mitochondrial phenotype are differentially responsive to growth of the female and the male fetus. This study may inform the design of sex-specific therapies for placental insufficiency and fetal growth abnormalities with life-long benefits for the offspring.

## Introduction

A successful pregnancy strongly depends on balancing resource allocation between the genetically determined fetal drive for growth and the mother who needs resources to support the pregnancy state. As the functional interface between mother and fetus, the placenta plays a key role in balancing fetal and maternal resource needs. Amongst its plethora of functions, the placenta executes the metabolism and secretion of hormones that have physiological effects on the mother and fetus, and transfers nutrients and oxygen from the mother to the fetus [1]. Thus it is perhaps unsurprising that fetal weight is related to placental development, the uteroplacental blood supply of nutrients and oxygen, and the capacity of the placenta to transport substrates to the fetus [2–8]. Moreover, failure of the placenta to grow and function properly is associated with the divergence of the fetus away from their genetic growth potential and can lead to small for gestational age (SGA), fetal growth restriction (FGR) or large for gestational age (LGA) [9–11]. SGA, FGR and LGA not only increase the risk of perinatal morbidity and mortality, but also have long-term consequences for offspring health [12]. Thus, it is important to understand the placental mechanisms regulating fetal growth outcomes.

To enable normal placental grow and function, the placenta depends on energy supplied by mitochondria. Mitochondria are the primary source of ATP, which is produced by oxidative phosphorylation (OXPHOS) using substrates derived from β-oxidation and the tricarboxylic acid cycle. ATP is used by the placenta to fuel growth and placental endocrine and transport functions. Mitochondria are also the place within the cell where steroidogenesis occurs; they contain several key proteins and enzymes such as STAR and CYP11A1, which are required for glucocorticoid and sex-steroid synthesis [13]. Mitochondria are also involved in cell signalling, homeostasis and survival via production of reactive oxygen species (ROS) and other molecules like nitric oxide. They are also dynamic organelles that can replicate (biogenesis), divide (fission) and combine (fusion) in response to metabolic, growth and stress signals [14, 15]. During pregnancy, there are temporal changes in placental mitochondrial respiratory capacity and mitochondrial-related proteins in several species [14,16–19].

Increasing evidence also suggests that placental mitochondrial function (mitochondrial OXPHOS, abundance, biogenesis, fission-fusion and efficiency) alters in line with defects in fetal growth and placental development in response to experimental reductions maternal nutrient and oxygen availability [20–23]. However, little is known about the relationship between placental mitochondrial capacity, placental morphological development and natural deviations in fetal growth in normal, uncompromised pregnancies. Even less is known about whether this relationship may vary for female and male fetuses, which is highly relevant given that sex is emerging as an important contributor to changes in placental, fetal and offspring health outcomes [24, 25].

In this study, we employed an integrative approach to evaluate placental morphology, mitochondrial OXPHOS capacity and mitochondrial regulator expression (electron transport system (ETS) complexes and biogenesis and fission-fusion regulators), in relation to growth of the lightest and heaviest female and male fetuses within the litter of normal wildtype mice. Importantly, since the mouse is a polytocous species, normal variation of fetal weight is expected within the litter, even under a normal, healthy gestational environment. We also examined the activity of signalling pathways governing placental growth and metabolism, as well as the expression of nutrient transporter and steroid hormone handling genes to further understand how placental phenotype is modulated by fetal weight and sex within the litter. Analyses were conducted on the labyrinth zone (Lz) of the mouse placenta as it is responsible for controlling the transport of nutrients, oxygen and hormones from mother to fetus

## Methods

### Animals

All experiments were performed under the U.K. Animals (Scientific Procedures) Act 1986 after ethical approval by the University of Cambridge. A total of 12 C57BL/6J virgin female mice were housed in the University of Cambridge Animal Facility using a 12/12 dark/light system and received *ad libitum* water and chow food (Rodent No. 3 breeding chow; Special Diet Services, Witham) during the study. At 4 months of age, females were mated overnight and the day a copulatory plug was found was designated as gestational day (GD) 1. On GD18, pregnant dams were killed by cervical dislocation, uteri were recovered, and fetuses and placentas cleaned from fetal membranes. All fetuses and placentas from the litter were weighed. For 7 litters, the placental Lz was micro-dissected, to separate it from the placental endocrine junctional zone and maternal decidua[26]. Lz samples were then cut in half, and one Lz half was placed into ice-cold biopsy preservation medium (0.21 M Mannitol, 0.07 M Sucrose, 30% DMSO in H_2_O and pH7.5) and frozen at -80°C until respirometry analysis. The remaining half of the Lz was snap frozen and stored at -80°C for molecular analyses (RT-qPCR and Western Blot). Placental samples from the additional 5 litters were kept whole, placed into 4% paraformaldehyde, histologically processed, and used for structural analysis. Fetal brain and liver were dissected and weighed from each fetus. Fetal tails were kept for sex determination by detection of the *Sry* gene using the Taq Ready PCR system (Sigma), specific primers (*Sry*: FPrimer: 5’-GTGGGTTCCTGTCCCACTGC-3’, RPrimer: 5’- GGCCATGTCAAGCGCCCCAT-3’ and PCR autosomal gene control: FPrimer: 5’- TGGTTGGCATTTTATCCCTAGAAC-3’, RPrimer: 5’-GCAACATGGCAACTGGAAACA-3’) and agarose gel electrophoresis.

### Placental Lz mitochondrial respiratory capacity

High resolution respirometry (Oxygraph 2k respirometer; Oroboros Instruments, Innsbruck, Austria) was used to assess the capacity for respiratory substrate use and ETS function. Cryopreserved Lz samples (10-15 mg) were gently thawed in ice-cold sucrose solution (0.25 M sucrose, 0.01 M TRIS-HCl, pH 7.5). Samples were then permeabilized in respiratory medium BIOPS (10 mM CaEGTA buffer, 0.1 µM free Ca^2+^, 1 mM free Mg^2+^, 20 mM imidazole, 20 mM taurine, 50 mM K-MES, 0.5 mM ditriothreitol, 6.56 mM MgCl2, 5.77 mM ATP and 15 mM phosphocreatine, pH 7.1) containing saponin (5 mg in 1 ml, Sigma-Aldrich, UK) for 20 minutes on ice. Samples were then washed by three 5 minute washes in respiratory medium MiR05 (0.5 mM EGTA, 3 mM MgCl_2_x6H_2_O, 20 mM taurine, 10 mM KH_2_PO_4_, 20 mM Hepes, 1 mg/mL of BSA, 60 mM K-lactobionate, 110 mM sucrose, pH 7.1) on ice to remove all endogenous substrates and contaminants. Oxygen concentration (µM) and flux per tissue mass (pmol O_2_/s/mg) were recorded in real-time using calibrated oxygen sensors and Datlab software (Oroboros Instruments, Austria). Respiratory rates were corrected for instrumental background by DatLab, considering oxygen consumption of the oxygen sensor and oxygen diffusion out of or into the oxygraph chamber measured under experimental conditions in miR05 medium without any tissue present.

A substrate-inhibitor titration protocol was performed under the presence of octanoylcarnitine and using approximately 10-15 mg of permeabilized Lz tissue placed into each oxygraph chamber. Briefly, complex I substrate malate (2mM) was added first to determine LEAK respiration (uncoupled from ATP synthesis; complex I LEAK or CI_Leak_). Next, ADP (5 mM), pyruvate (20mM) and glutamate (10 mM) were added to obtain complex I oxygen flux under OXPHOS state (CI_P_). Then, succinate (10 mM) was added to provoke complex I and II dependent oxidative phosphorylation (CI+ÍI_P_). Fatty acid oxidation (FAO) was calculated immediately after ADP addition (i.e. octanoylcarnitine, plus malate and ADP). Trifluoromethoxy carbonyl- cyanide phenylhydrazone (FCCP, two doses of 0.5 µM each) was added to obtain total ETS capacity (uncoupled state). To activate complex IV (CIV) dependent respiration, the first three complexes of the ETS were inhibited by adding rotenone (inhibits complex-I; 0.5 µM), malonic acid and myxothiazol (inhibits complex-II; 5 mM and 0.5 µM, respectively) and antimycin A (inhibits complex-III; 2.5 µM). Sodium ascorbate (2 mM) and N, N, N’, N’-tetramethyl-p-phenylenediamine (TMPD, 0.5 mM) were then added to stimulate complex IV supported respiration, which was then inhibited by adding sodium azide (200 mM). ETS excess capacity was calculated using the formula: 1- CI+ÍI_P_/total ETS (1-P/E). Cytochrome c (10 µM) was added to check mitochondrial membrane integrity and data were excluded if respiration increased by >30%. All substrates used were at their saturating concentrations to assess maximal mitochondrial respiratory capacity.

### Placental Lz gene expression analysis

Placental Lz RNA was extracted using the RNeasy Plus Mini Kit (Qiagen, Hilden, UK) and the quantity of RNA obtained was determined using a NanoDrop spectrophotometer (NanoDrop Technologies, Inc., Auburn, AL). A total of 2 μg per sample was reverse transcribed using the High Capacity cDNA Reverse Transcription Kit (Applied Biosystems, Foster City, USA) according to the manufacturer’s instructions. Three dilutions of each cDNA sample (1:10, 1:20 and 1:100) were run as a triplicate along with non-template controls in the 7500 Fast Real-Time PCR thermocycler System (Applied Biosystems, UK) for gene expression quantification using gene specific primer pairs (Table 1) and SYBR Green master-mix (Applied Biosystems, UK). The standard thermal cycling protocol was conducted as follows: 50 °C for 2 min, 95 °C for 10 min and 40 cycles of 95 °C for 95 seconds and 60 °C for 1 min. Relative expression was calculated using the 2^-ΔΔCt^ method and genes of interest were normalized to the mean expression of 3 housekeeping genes (*Hprt*, *Ywhaz* and *Ubc*), which were stable in the placental Lz between the lightest and heaviest fetuses of each sex. Data were then displayed relative to the average mRNA expression value for the heaviest fetus of each sex.

**Table 1.**
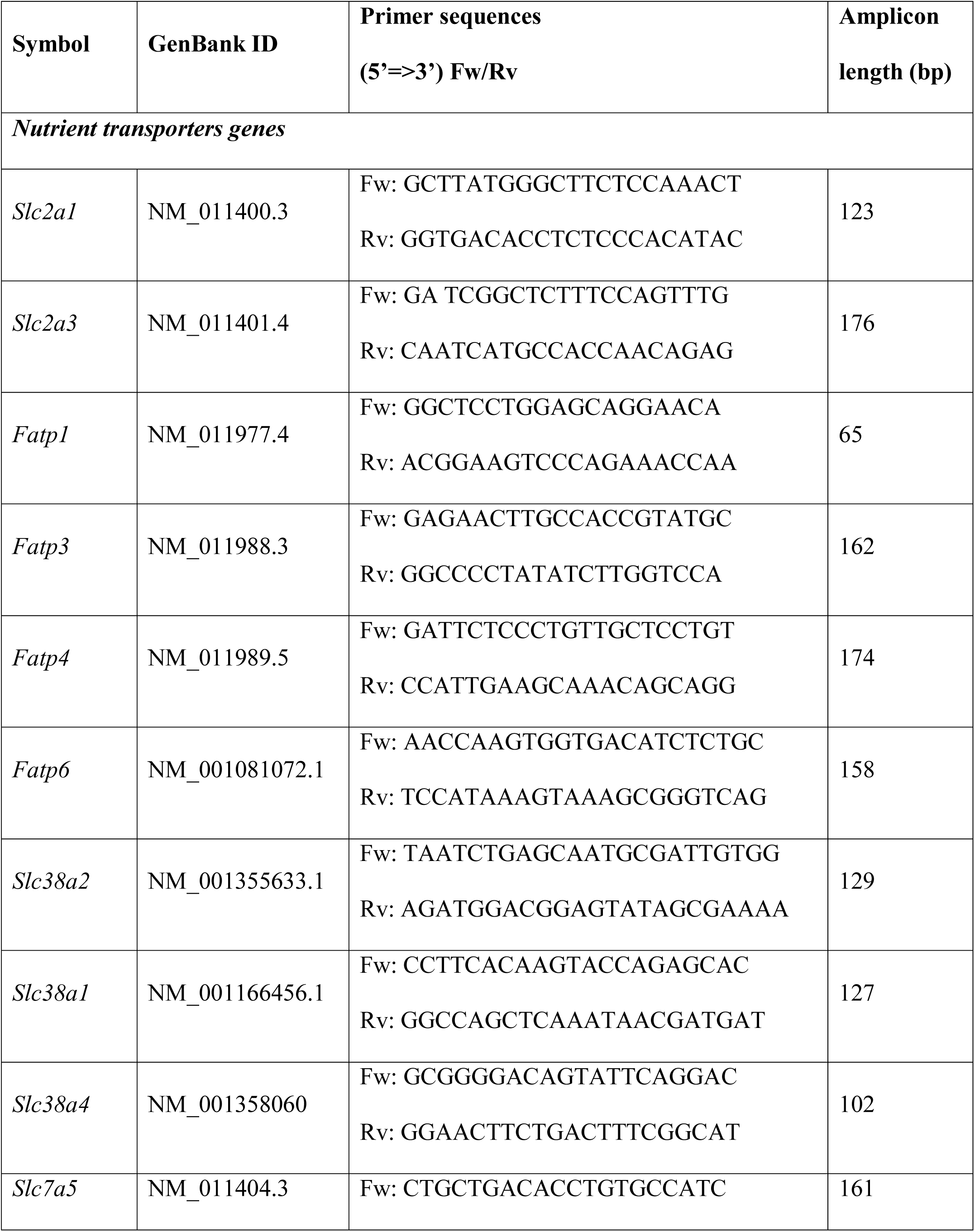
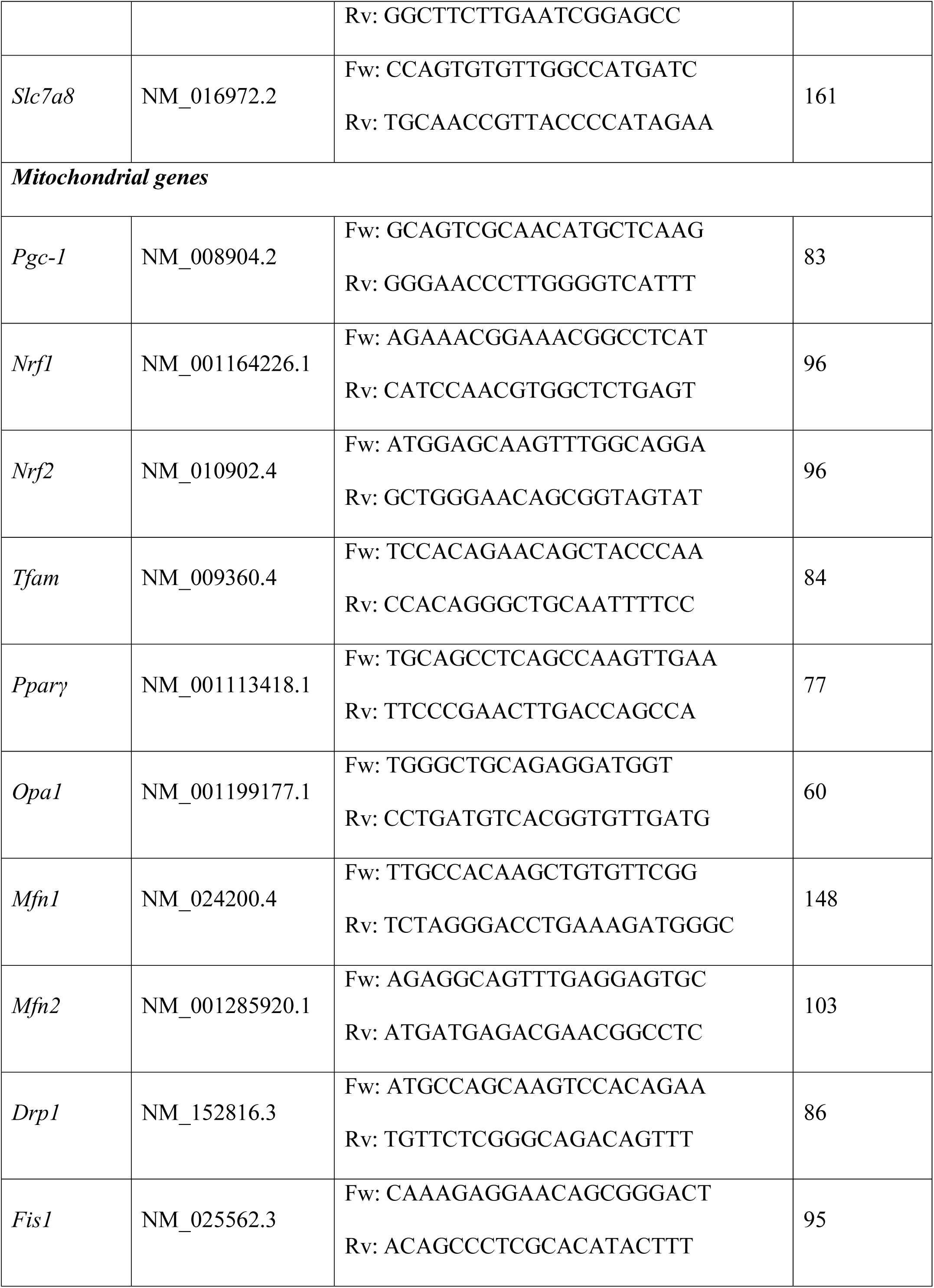
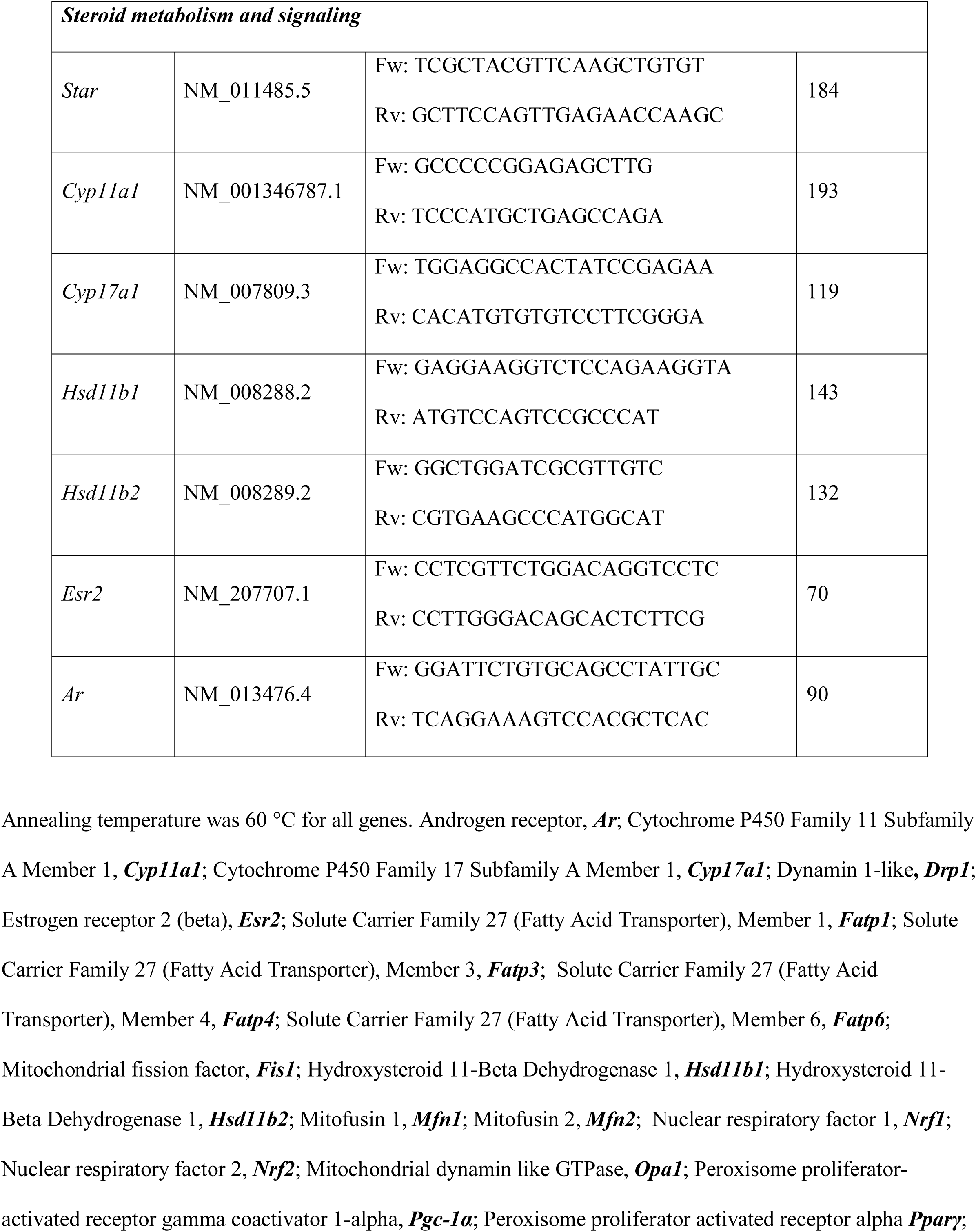
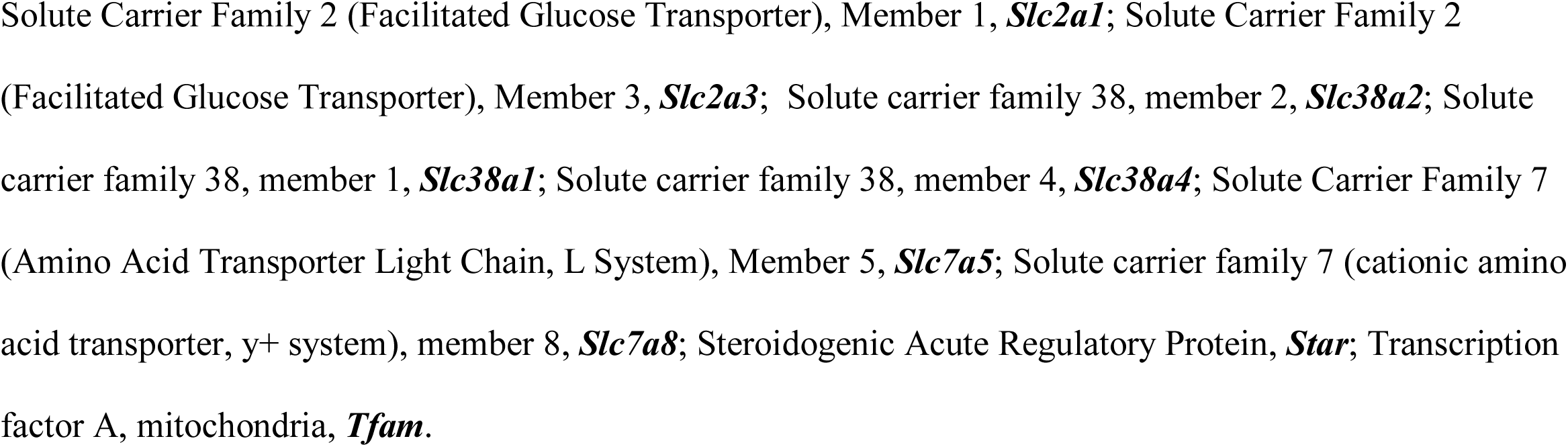
Primers used for qPCR analysis.

### Placental Lz protein abundance analysis

Protein extraction was performed on frozen Lz tissue homogenized using commercial RIPA lysis buffer (Thermo Scientific, US) supplemented with Mini EDTA-free protease inhibitor cocktail mix (Roche, CH). The protein concentration was determined using the Bicinchoninic Acid protein assay (Thermo Scientific, US). Lysates (3μg/μl in 1xSDS) were separated by sodium dodecyl sulphate polyacrylamide gel electrophoresis (SDS-PAGE) and transferred onto 0.2 μm nitrocellulose membranes (Bio-Rad Laboratories, US) using a semi-dry technique (Semi-dry Blotter, Invitrogen). Membranes were stained in Ponceau red and staining captured on an iBRIGHT (Thermo Scientific, US). The membrane was washed with tris-buffered saline tween (TBST) and blocked with 5% milk or fetal bovine serum (used for phosphorylated proteins) in TBST on a shaker, for 60 minutes at room temperature. Membranes were then incubated overnight at 4°C with primary antibody (Table 2). The day after, membranes were washed and incubated with rabbit or mouse secondary antibodies tagged to horseradish peroxidase (NA934 or NA931; 1:10000) diluted in TBST containing 2.5% milk for 60 minutes. Protein bands were visualised using Scientific SuperSignal West Femto enhanced chemiluminescence (ECL) substrate (Thermo Scientific, US) and captured on an iBRIGHT. The signal intensity of protein bands were quantified using ImageJ software and normalized for slight variations in loading using a corresponding band of similar molecular weight on the captured Ponceau red stained membrane [27]. The abundance of phosphorylated proteins was calculated as a ratio to their respective total protein abundance. Data are displayed relative to the average protein expression value for the heaviest fetus of each sex.

**Table 2.**
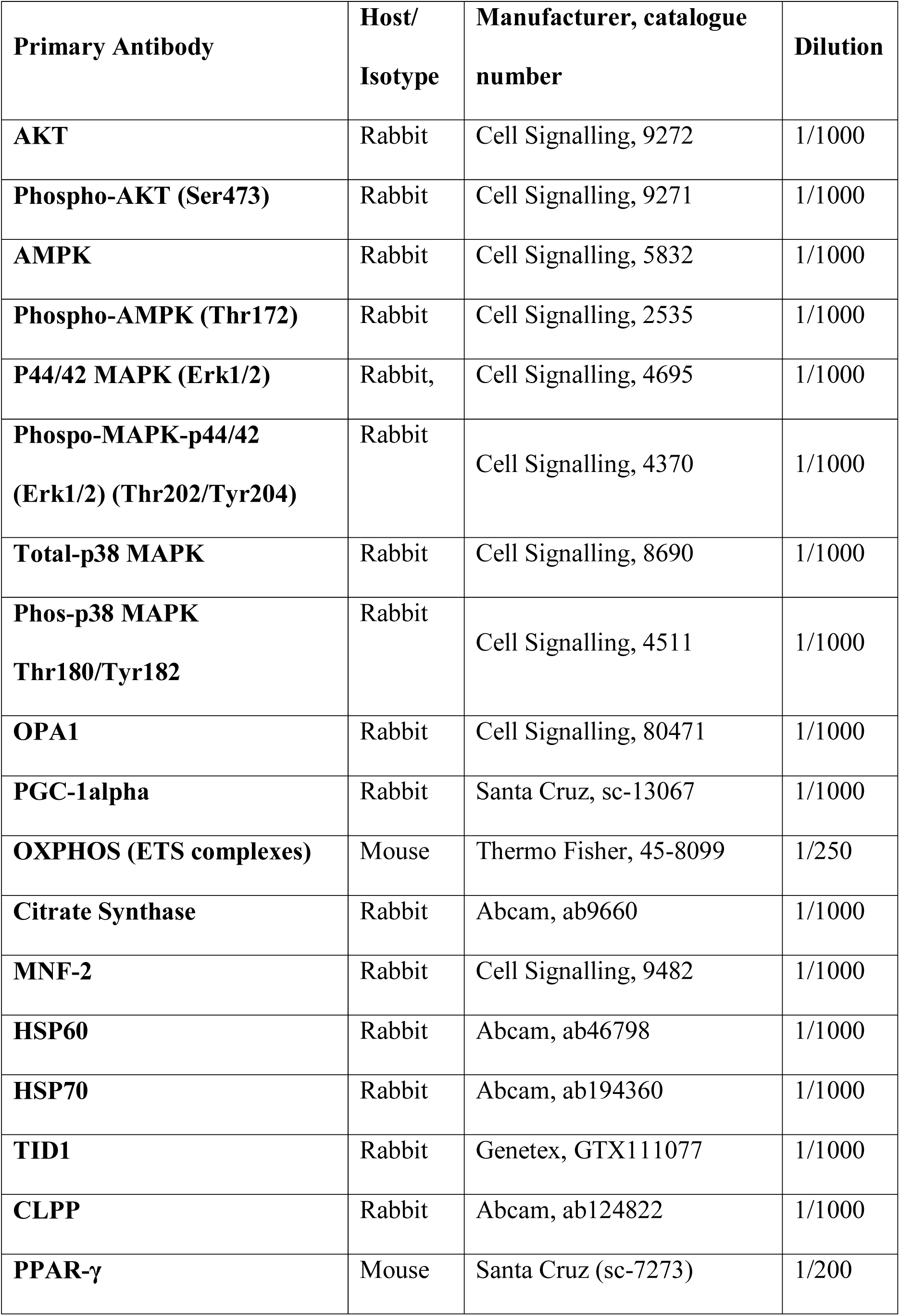
List of primary antibodies used in this study.

### Placental Lz structural analysis

Morphology of the Lz was assessed by double-labelling placental sections with cytokeratin and lectin antibodies to identify trophoblast and fetal capillaries, respectively. Details about the staining protocol have been described in detail elsewhere [28]. Stained sections were then scanned using a NanoZoomer 2.0-RS Digital Pathology System (NDP Scan Hamamatsu, Japan) and stereological analysis of the Lz was performed as described previously [28].

### Statistical analysis

Statistical analyses were performed using GraphPad Prism version 8 (GraphPad, La Jolla, CA, USA). To check whether each data set was normally distributed, Shapiro-Wilk tests were performed, and identified that all displayed a normal Gaussian distribution. To compare conceptus biometry between males and females within the litter, paired *t*-tests were performed, allowing the elimination of inter-litter variation. Similarly, paired *t-*tests were performed to assess differences in placental structure, mitochondrial respiration, and gene and protein expression between the heaviest and the lightest fetuses of each sex within the litter Retrospective comparisons between the lightest females and lightest males and heaviest females and heaviest males within the litter were also performed by paired *t-*tests. Relationships between data were undertaken using Pearson’s correlation coefficient (r). Values are expressed as individual data points and/or mean ± SEM and p values < 0.05 were considered statistically significant. The number of samples per group for each analysis is shown in each figure and described in the legends of figures and footnotes of tables.

## Results

### Conceptus biometry for all female *versus* all males within the litter

Considering all fetuses together, conceptus biometric data were different between females and males within the litter at gestational day 18 (term on day 20; Figure 1). In particular, fetus, placenta and Lz weights were lower in females when compared with male fetuses (p=0.03, p<0.001 and p=0.02, respectively). However, there were no differences in fetal brain and liver weights (relative to body weight), or placental efficiency (p=0.06), calculated as the ratio of fetal weight to placental weight between females and males (Figure 1).

**Figure 1.**
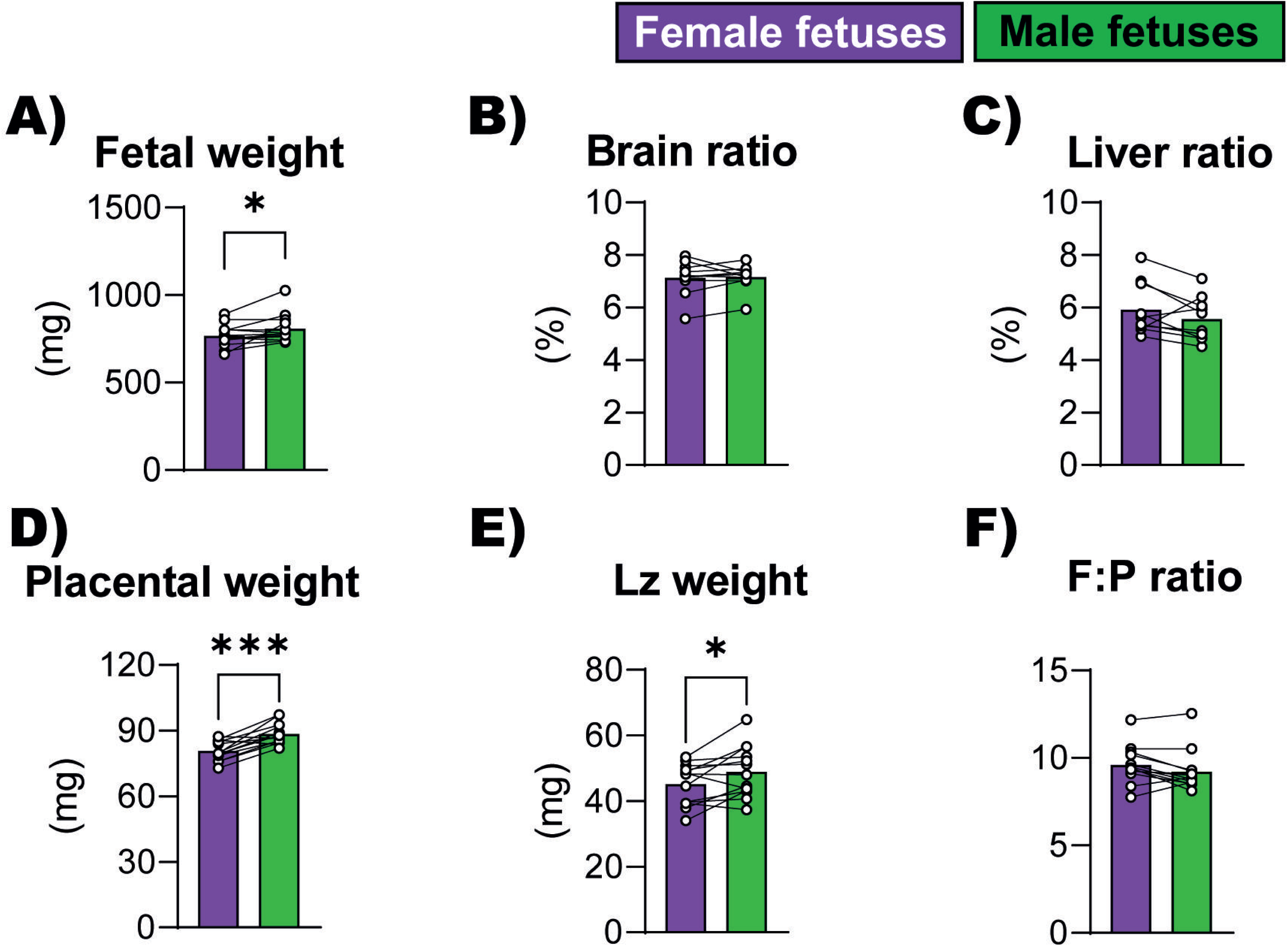
Conceptus biometrical data from female and male fetuses within the same litter at gestational day 18. (A) Fetal weight, (B) Brain and (C) Liver weights as a proportion of fetal weight, (D) Placental weight, (E) Labyrinth zone (Lz) weight and (F) Placental efficiency (F:P ratio; determined as the ratio of fetal weight to placental weight). Data are from n = 12 litters were averaged. Data are displayed as individual litter data points with bar representing the mean value and lines connecting siblings from the same litter. Data analysed by paired t test; ∗p < 0.05, ∗∗∗p < 0.001.

Placental weight was not significantly correlated with fetal weight when analysing all conceptuses within the litter collectively (n=71; r=0.17; p=0.15). Similarly, when data was separated into females and males, again no correlation was detected (Figure 2A, females: n=35; r=0.24; p=0.15, males: n=36; r=-0.07 p=0.64). However, when data were segregated to only assess the heaviest and lightest fetuses per sex within the litter, a positive correlation between placental and fetal weight was found for the lightest females (Figure 2B, r=0.74; p=0.005), but not for the lightest males (Figure 2C, r=0.32; p=0.31) or the heaviest fetuses of either sex. These data suggest that the placenta may be supporting growth of the female and male fetuses in different ways within the litter.

**Figure 2.**
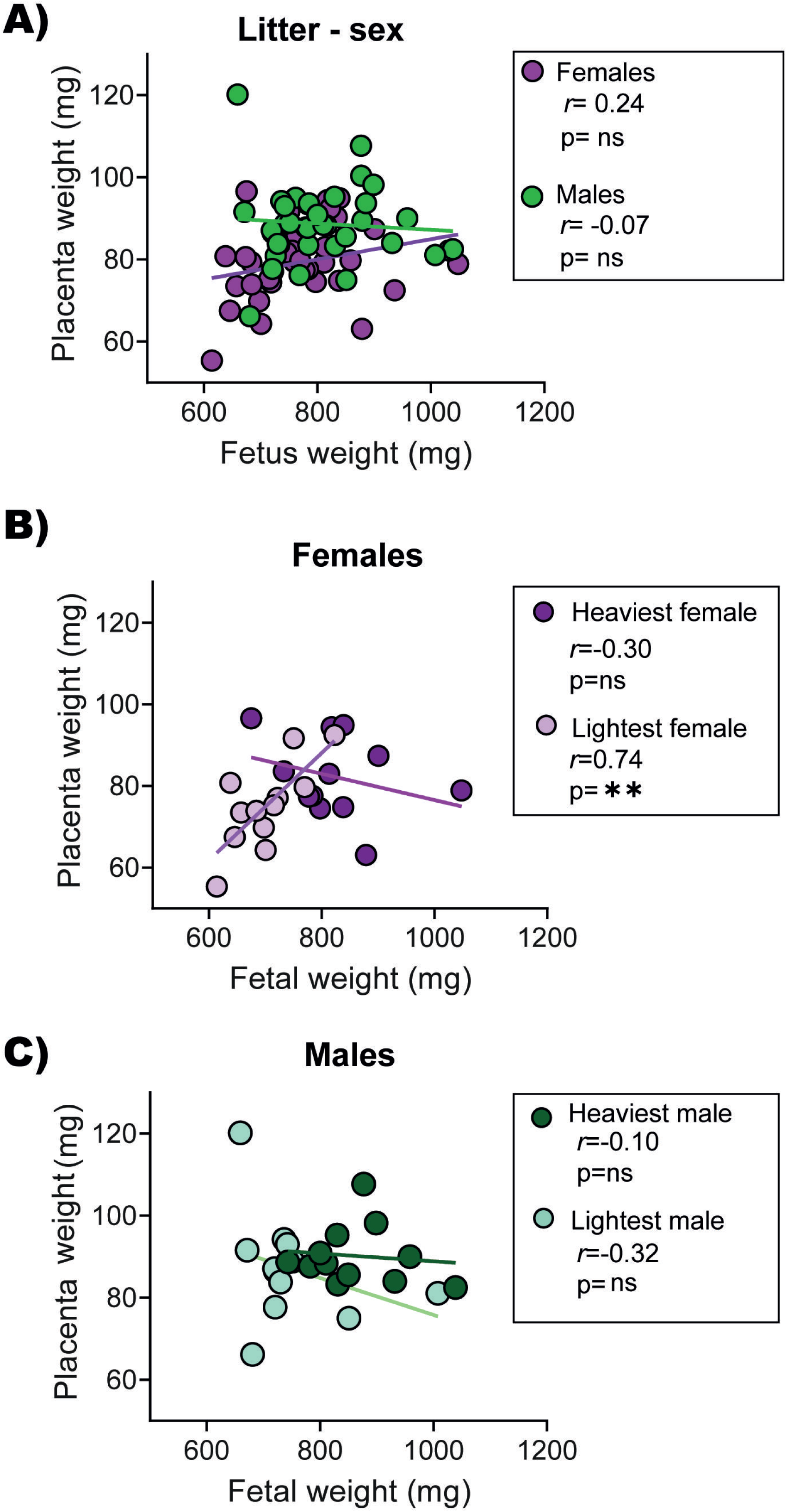
Relationships between placental and fetal weights (A) for each fetal sex when using all data obtained for each litter, (B) using data obtained for just the lightest and heaviest female fetuses in the litter, and (C) using data obtained for just the lightest and heaviest male fetuses in the litter on gestational day 18. Data were analysed using Pearson’s correlation coefficient and *r* values are shown in each corresponding figure (∗∗p < 0.01). In A, data are from 35 females and 36 males from n = 12 litters. In B and C, data are from 12 lightest and 12 heaviest fetuses and respective placentas per sex from n = 12 litters. ns = not significant.

### Conceptus biometry and placental Lz morphology for the lightest *versus* the heaviest fetuses of each sex

To understand why there may be a sex-related difference in the relationship between placental weight and fetal weight, the lightest and heaviest fetuses from each litter were selected and conceptus biometry were compared for each sex separately (Figure 3). As expected, fetal weight was lower for the lightest compared to the heaviest for each fetal sex in the litter (Figure 3A, females; p=0.002; males; p<0.0001), and the mean weight difference between them were similar for females and males (14.1% and 13.6% less than heaviest, respectively). Fetal brain and liver weights as a proportion of body weight did not vary, which suggests that the lightest fetuses are symmetrically smaller when compared to the heaviest fetuses (Figure 3B-D). Moreover, placental weight, Lz weight and placenta and Lz efficiency did not vary between the lightest and the heaviest fetuses within the litter, regardless of fetal sex (Figure 3E-H)

**Figure 3.**
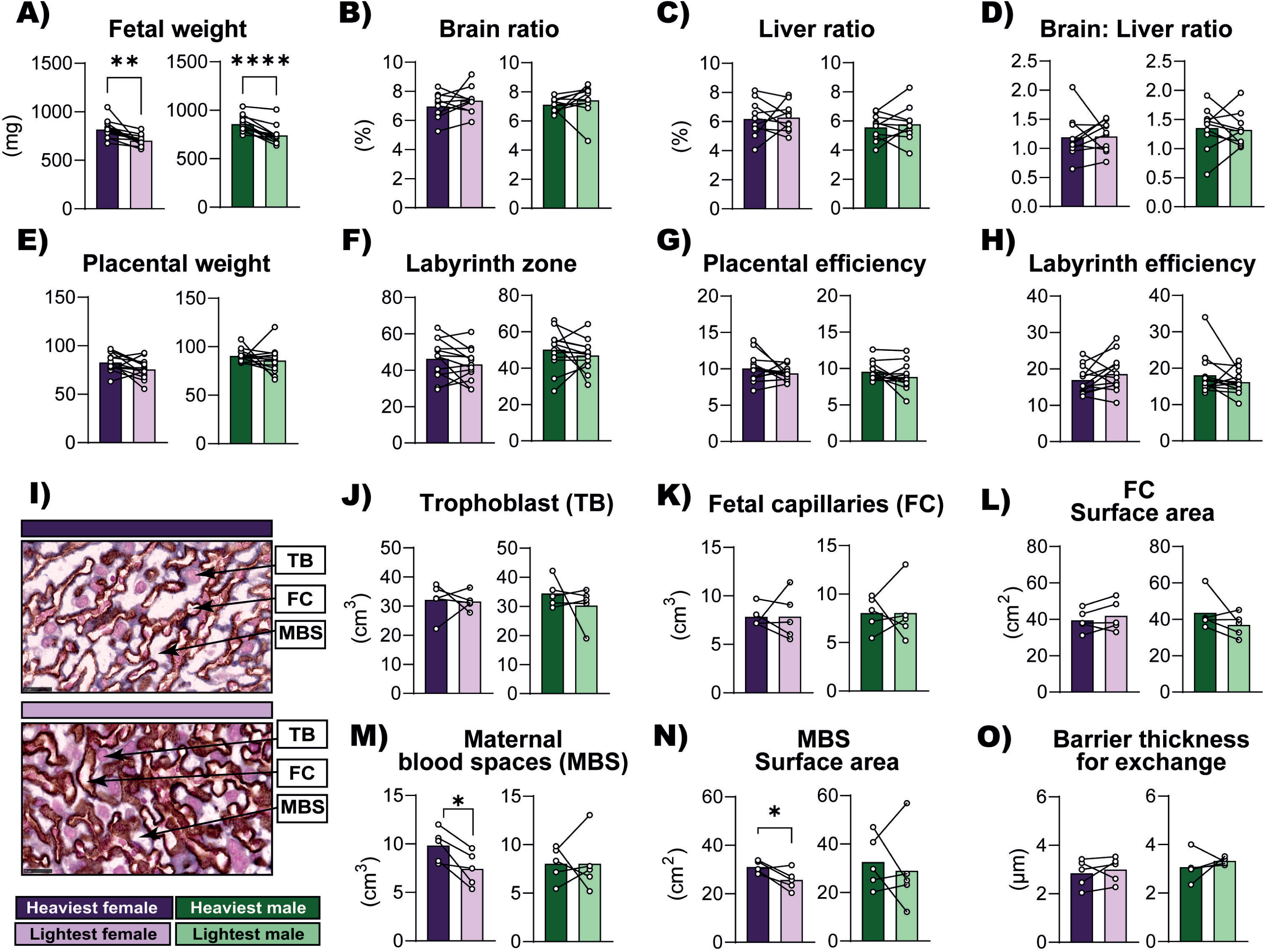
Conceptus biometry and labyrinth zone structure of the lightest *versus* the heaviest fetuses of each fetal sex within the litter on gestational day 18. (A) Fetal weight (B) Brain and (C) Liver weights as a proportion of fetal weight, (D) Brain weight to liver weight ratio, (E) Placenta weight, (F) Labyrinth zone weight, (G) Placental efficiency, (H) Labyrinth efficiency (determined as the ratio of fetal weight to labyrinth weight), (I) Representative images of placental labyrinth histology of female fetuses, (J) Trophoblast volume, (K) Fetal capillaries volume, (L) Fetal capillaries surface area, (M) Maternal blood spaces volume, (N) Maternal blood spaces surface area, and (O) Barrier thickness. For conceptus biometry analysis, 12 lightest and 12 heaviest fetuses and respective placentas per sex were used. For stereological analysis, 5 lightest and 5 heaviest fetuses and respective placentas per sex were used. Data are displayed as individual data points with bar representing the mean value and lines connecting siblings from the same litter. Data analysed for each sex separately by paired t test; *p < 0.05 ∗∗p < 0.01 and ∗∗∗∗p < 0.0001. Abbreviations: TB (trophoblasts); FC (Fetal capillaries); MBS (Maternal blood spaces)

Stereological analysis of the placental Lz zone revealed that there were no differences in trophoblast and fetal capillary volumes (Figure 3I-K). However, there was less maternal blood spaces in the placental Lz of the lightest females, compared to the heaviest females, and this difference was not found for the males (Figure 3M). Similarly, maternal blood space surface area was lower in the lightest, compared to heaviest female fetuses, an effect not observed for the male fetuses (Figure 3N). The surface area of the fetal capillaries (Figure 3L) and barrier thickness (Figure 3O) of the Lz did not vary between the lightest and the heaviest fetuses, for either females or males.

### Mitochondria respiratory capacity of the placental Lz for the lightest *versus* the heaviest fetuses of each sex

To investigate whether sex-dependent structural changes in the Lz zone between the lightest and heaviest fetuses may be related to mitochondrial functional alterations, high resolution respirometry was performed (Figure 4A). Oxygen flux rate analysis revealed that in LEAK state, mitochondrial CI related oxygen consumption was ∼60% greater for the placental Lz of lightest compared to the heaviest females, but no effect was seen for the males (Figure 4B, p=0.003). Whilst CI oxygen flux under OXPHOS state was not different between the lightest and heaviest fetuses of either sex (Figure 4C), after adding succinate, Lz CI+CII oxygen consumption rate was ∼44% greater in the lightest compared to the heaviest females within the litter (p=0.01); a difference that was not observed for the males (Figure 4D). Fatty acid oxidation (FAO), total ETS capacity and CIV associated oxygen consumption rates by the placental Lz were not different between lightest and heaviest fetuses for either fetal sex (Figure 4E-G). When oxygen consumption rates for CI in LEAK state and CI+II in OXPHOS state were corrected to total ETS oxygen flux to provide a qualitative indication of changes in mitochondrial function per mitochondrial unit, these values were also increased in only the lightest compared to heaviest females (not different for the lightest compared to heaviest males) (Figure 4H, p=0.02; 3I and 3J, p=0.03). In addition, calculation of 1-P/E indicated that ETS excess capacity was lower in the lightest females compared to the heaviest (p=0.03), again a difference not identified for the males (Figure 4K).

**Figure 4.**
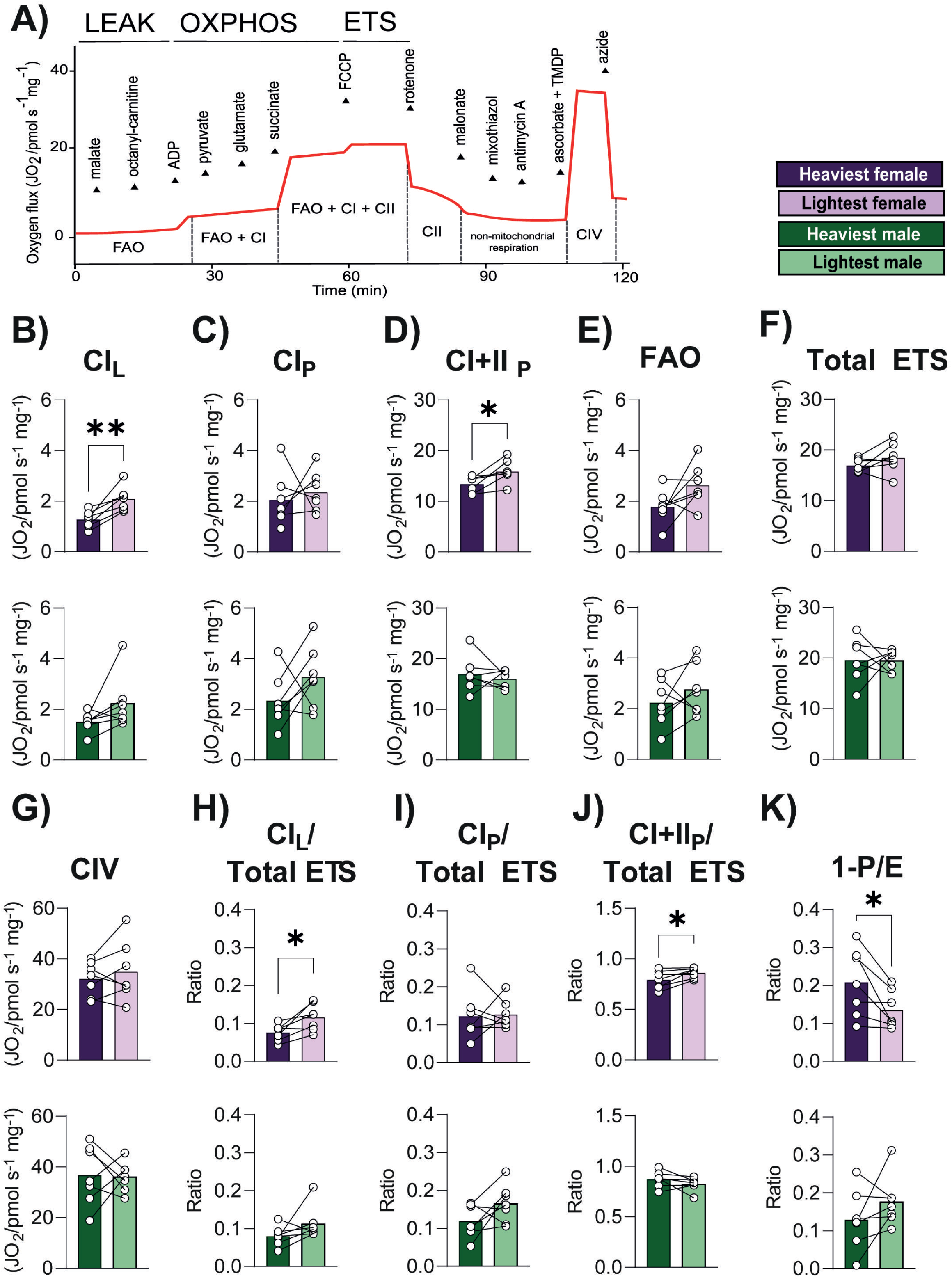
Mitochondrial respiration rates in the labyrinth zone supporting the lightest and heaviest fetuses of each sex within the litter on gestational day 18. (A) Representative experimental trace, (B) C_L_: C_Leak_, (C) C_P_: C_Oxphos_, (D) CI+II_P_: CI+CII_Oxphos_, (E) FAO, (F) Total ETS, (G) CIV, (H) C_L_/ Total ETS, (I) C_P_/ Total ETS, (J) CI+II_P_/ Total ETS and (K) 1-P/E. Sample size was 7 lightest and 7 heaviest fetuses per sex from n = 7 litters. Data are displayed as individual data points with bar representing the mean value and lines connecting siblings from the same litter. Data analysed for each sex separately by paired t test; ∗p < 0.05 and ∗∗p < 0.01.

### Expression of mitochondrial ETS components, dynamic genes, and regulatory proteins in the placental Lz for the lightest *versus* the heaviest fetuses of each sex

To gain further information on the sex-dependent differences in placental mitochondrial respiratory capacity, western blotting and qPCR was performed to determine the expression ETS complex proteins (CI-V), biogenesis, fusion and fission genes, and additional mitochondrial regulatory proteins in the placental Lz of the lightest and heaviest fetuses for both sexes (Figure 5). These analyses revealed that CI protein abundance was lower in the lightest compared to heaviest females (Figure 5A), meanwhile CIII and CV proteins were lower only in the lightest compared to the heaviest males (Figure 5B). In addition, the expression of mitochondria biogenesis gene, *Nrf1* and mitochondrial fission genes, *Drp1* and *Fis1,* was lower in the Lz of the lightest females, when comparing with the heaviest females (Figure 5C). Whereas the expression of *Tfam,* a mitochondria biogenesis transcription factor gene, was greater in the Lz of the lightest males versus the heaviest males (Figure 5D). Mitochondrial content, informed by citrate synthase protein abundance, did not vary in the Lz between the lightest and heaviest fetuses within the litter, regardless of fetal sex (Figure 5E). In addition, abundance of mitochondrial biogenesis (PGC-1α), fusion (MNF2 and OPA1), heat shock (HSP60, HSP70) and chaperone (TID1) proteins did not differ in the Lz between the lightest and heaviest fetuses within the litter, in either sex (Figures 5E). However, protein abundance of CLPP a key protease involved in mitochondrial protein clearance and a marker of the mitochondrial unfolded protein response (UPRmt), was lower in the lightest females compared with the heaviest females; an effect not seen for males (Figure 5E).

**Figure 5.**
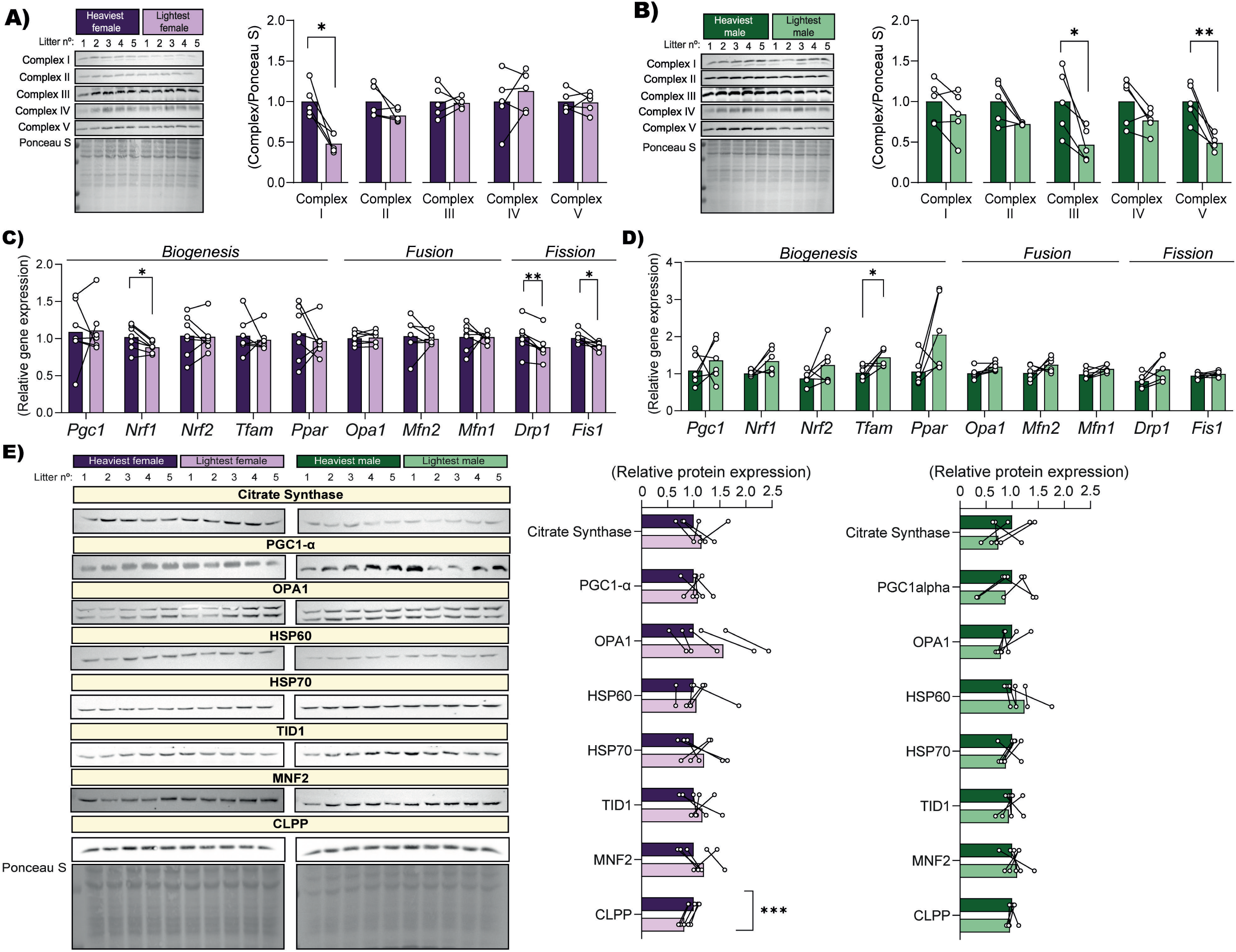
Protein abundance of electron transport chain complexes (A and B, males and females, respectively). Relative mRNA expression of key mitochondria dynamics genes associated with biogenesis, fusion, and fission processes on females (C) and males (D) and key mitochondrial regulatory proteins; (E) citrate synthase, PGC1α MNF2, OPA1, HSP60, HSP70, TID1 and CLPP in the placental labyrinth zone supporting the lightest and heaviest fetuses of each sex within the litter on gestational day 18. Images from each antibody and representative Ponceau staining are included. For western blot results sample size is 5 lightest and 5 heaviest fetuses per sex from n = 5 litters. For qPCR results sample size is 7 lightest and 7 heaviest fetuses per sex from n = 7 litters. Data are displayed as individual data points with bar representing the mean value and lines connecting siblings from the same litter. Data are displayed relative to the value for the heaviest fetus per sex. Data were analysed for each sex separately by paired t test; ∗p < 0.05, ∗∗p <0.01, ∗∗∗p <0.001.

### Expression of transport genes and steroid metabolism and signalling genes in the placental Lz for the lightest *versus* the heaviest fetuses of each sex

Since the energy provided by mitochondria helps to fuel placenta transport and endocrine function, we evaluated whether sex-related variations found on mitochondria functional capacity (respiratory function, gene and protein regulators) are associated with the expression of nutrient transporter and steroidogenic genes between the lightest and heaviest of each sex within the litter. In particular, the mRNA expression of key transporters for glucose (*Slc2a1* and *Slc2a3*), amino acid (*Slc38a1, Slc38a2, Slc38a4, Slc7a5* and *Slc3a2*) and lipids (*Fatp1, Fatp3, Fatp4, Fatp6* and *Cd36*) were quantified in the placental Lz zone by RT-qPCR (Figure 6). We also evaluated the expression of genes involved in steroid hormone production (*Star, Cyp11a1* and *Cy17a1*), glucocorticoid metabolism (*11bhsd1* and *11bhsd2*) and steroid hormone signalling (*Esr2* and *Ar*) in the placental Lz using qPCR. These analyses showed that the expression of *Slc2a1* mRNA was ∼20% lower for the lightest compared to the heaviest males (Figure 6B, p = 0.021), however this difference was not observed for the lightest *versus* the heaviest females. In addition, no differences were found between the lightest and the heaviest fetuses within the litter for any of the other nutrient transporter genes quantified in either fetal sex (Figure A-C). The gene expression of *Cyp11a1* was ∼63% greater (p=0.038), while *Cyp17a1* ∼20% lower (p=0.035) in the lightest compared to the heaviest female fetus, with no differences in these steroidogenic genes detected in the males (Figure 6D). Whereas the mRNA expression of the androgen receptor (*Ar*) a steroid-hormone activated transcription factor was ∼91% greater (p=0.046) in the lightest compared to the heaviest males only (Figure 6D). The mRNA expression of the estrogen receptor beta (*Esr2)*, the 11β-hydroxysteroid dehydrogenase, isozymes 1 and 2 (*Hsd11b1* and *Hsd11b2*) and the steroidogenic acute regulatory protein (*Star*) in the Lz were not different between the lightest compared to the heaviest fetuses in either sex.

**Figure 6.**
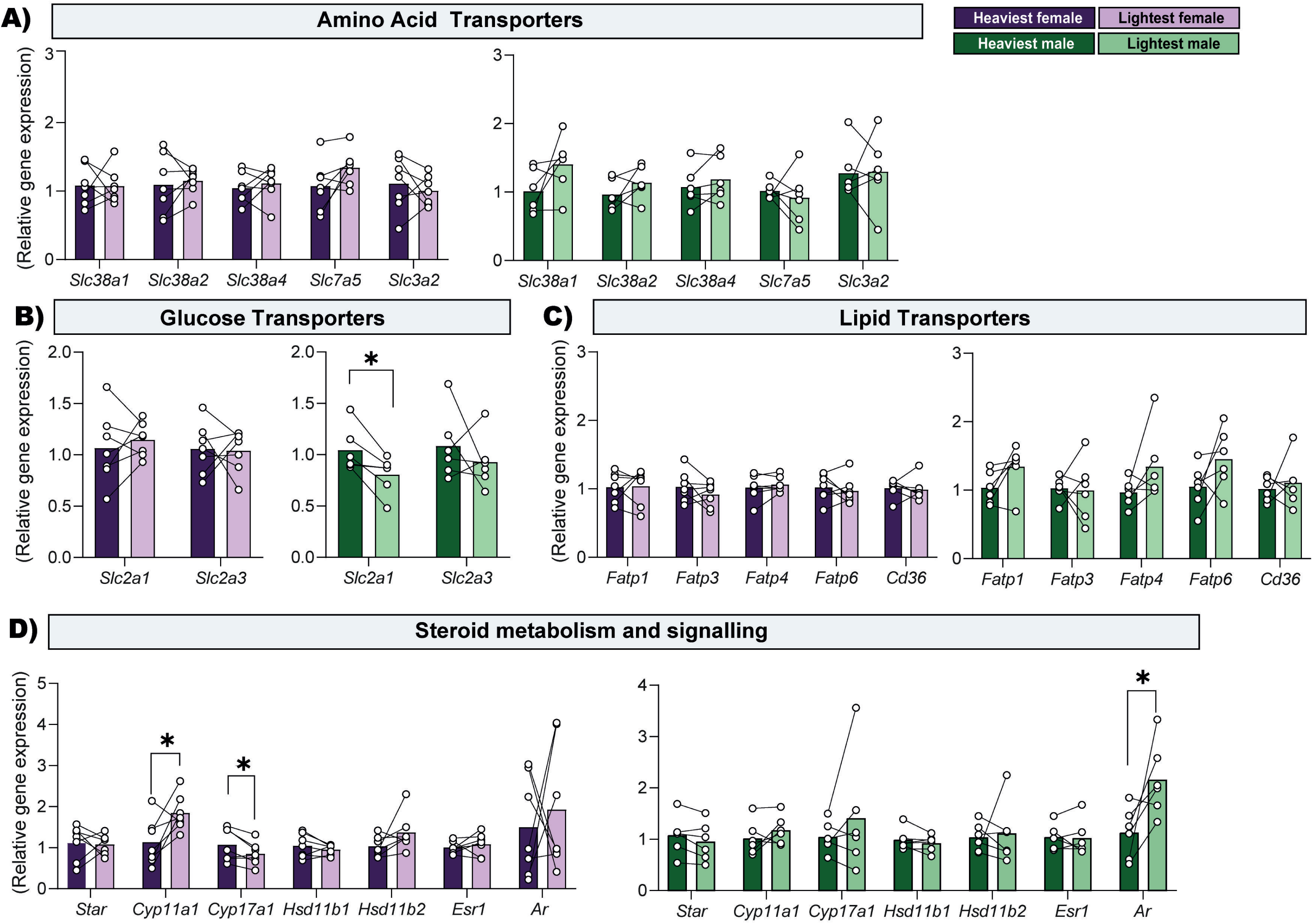
Relative mRNA expression of amino acid (A), glucose (B), lipid (C) transporters and steroid hormone metabolism and signalling related genes (D) in the placental labyrinth zone supporting the lightest and heaviest fetuses of each fetal sex within the litter on gestational day 18. Data are from 7 lightest and 7 heaviest fetuses per sex from n = 7 litters. Relative expression was calculated using the 2^-ΔΔCt^ method and genes of interest were normalized to the mean expression of 3 housekeeping genes (*Hprt, Ywhaz* and *Ubc*). Data are displayed as individual data points with bar representing the mean value and lines connecting siblings from the same litter. Data are displayed relative to the value for the heaviest fetus per sex. Data were analysed for each sex separately by paired t test; *p < 0.05.

### Abundance of key growth and metabolic proteins in the placental Lz for the lightest *versus* the heaviest fetuses of each sex

To provide information on the sex-dependent differences in placental morphology and mitochondrial function between lightest and heaviest fetuses, the abundance of key growth and metabolic signalling proteins, namely AKT (protein kinase B), AMPKα (5’-AMP-activated protein kinase catalytic subunit alpha-1), p44/42 MAP kinase (ERK1/ERK2), p38 MAPK (mitogen-activated protein kinase) and PPARγ (peroxisome proliferator-activated receptor gamma) were evaluated by western blotting (Figure 7). The abundance of total AMPKα protein was greater in the lightest compared to the heaviest fetuses for both females and males (67% and 41%, Figures 7A and C, p=0.005 and p=0.01, respectively), however this was not related to a significant change in activation of AMPKα (abundance of phosphorylated AMPKα normalized to total AMPKα protein, Figures 7B and D). While the total abundance of AKT protein did not vary between the lightest and the heaviest fetuses, activated AKT (phosphorylated to total AKT protein) was ∼32% lower in the Lz zone supporting the lightest compared to the heaviest males (p=0.032), but no difference was found for the females (Figures 7B and D). The abundance and activation of p44/42 MAPK and p38 MAPK proteins was not different between the lightest and the heaviest fetuses, irrespective of fetal sex (Figure 7A and D). Interestingly, abundance of PPARγ an important transcription factor involved mitochondrial metabolism and lipid synthesis, was **greater** in the lightest female compared to the heaviest female, whereas PPARγ protein was lower in the lightest males when compared to the heaviest males from the litter (Figure 7E, p=0.04 and p < 0.05, respectively).

**Figure 7.**
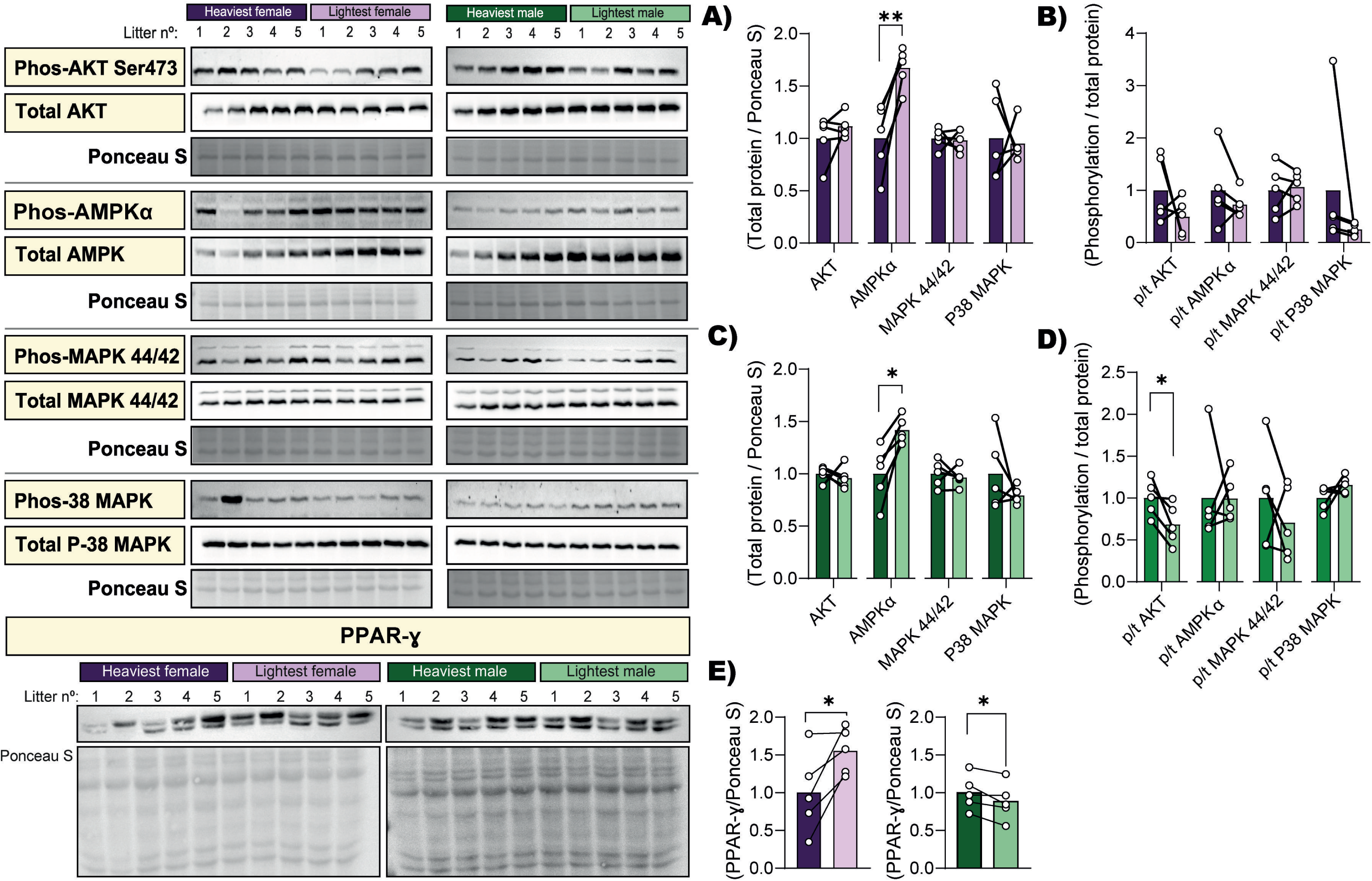
Protein abundance of key growth and metabolic signalling proteins in the placental labyrinth zone supporting the lightest and heaviest fetuses of each sex within the litter on gestational day 18. (A and C) Total AKT, AMPKα, MAPK 44/42 and P38 MAPK protein levels, and (B and D) AKT, AMPKα, MAPK 44/42 and P38 MAPK phosphorylation levels as a ratio to total protein in the heaviest *versus* the lightest fetuses for females (A and B) and males (C and D). Total protein abundance for PPARγ (E). Representative images from each antibody and Ponceau staining are included. Protein abundance and phosphorylation levels were normalized to Ponceau staining and total protein abundance, respectively. Data are 5 lightest and 5 heaviest fetuses per sex from n=5 litters. Data are displayed as individual data points with bar representing the mean value and the lines connecting siblings from the same litter. Data are displayed relative to the value for the heaviest fetus per sex. Data were analysed for each sex separately by paired t test; *p < 0.05

### Comparisons of the effect of sex on feto-placental growth of the lightest and heaviest fetuses

To gain further insight into the sex-dependent intra-litter differences in placental phenotype, retrospective comparisons between the lightest females and lightest males and heaviest females and heaviest males within the litter were performed (Table 3). These data showed that the heaviest males were ∼5% heavier than the heaviest female fetuses within the litter (p <0.05). The placenta of the heaviest males in the litter was also greater by ∼13% when compared to the heaviest females (p=0.04), with a tendency for this to also vary with sex for the lightest female littermates (p=0.054). The placental expression of glucose (*Slc2a1*: -37%, p=0.003) and lipid (*Fatp1*: +17%, tendency p=0.07) transporter genes were also differentially expressed between the lightest (but not heaviest) male and female fetuses of the litter. Placental respirometry rates associated with CI_Oxphos_ (+39%, tendency p=0.05) and with CI+CII_Oxphos_/Total ETS (+30%, tendency p=0.08) together with biogenesis (*Nrf1*: +37%, p=0.03; *Tfam*: +25%, tendency p=0.06) and dynamic (*Opa1*: +13%, tendency p=0.06, *Mfn1*: +11%, p=0.05) genes were all greater in the lightest males compared to the lightest females. Meanwhile, biogenesis genes *Pparγ* (-38%, p=0.03) and *Tfam* (-19%, tendency p=0.07) were decreased in the heaviest males compared to females. Finally, the expression of the steroidogenic gene *Cyp11a1* was lower (-38%, p=0.02) in the lightest males compared to females and in heaviest fetuses, gene expression of *Cyp17a1* (-41%, p=0.01) were decreased on males compared to females. There was also no effect of fetal sex in the placental Lz morphology of the lightest and heaviest fetuses.

**Table 3.**
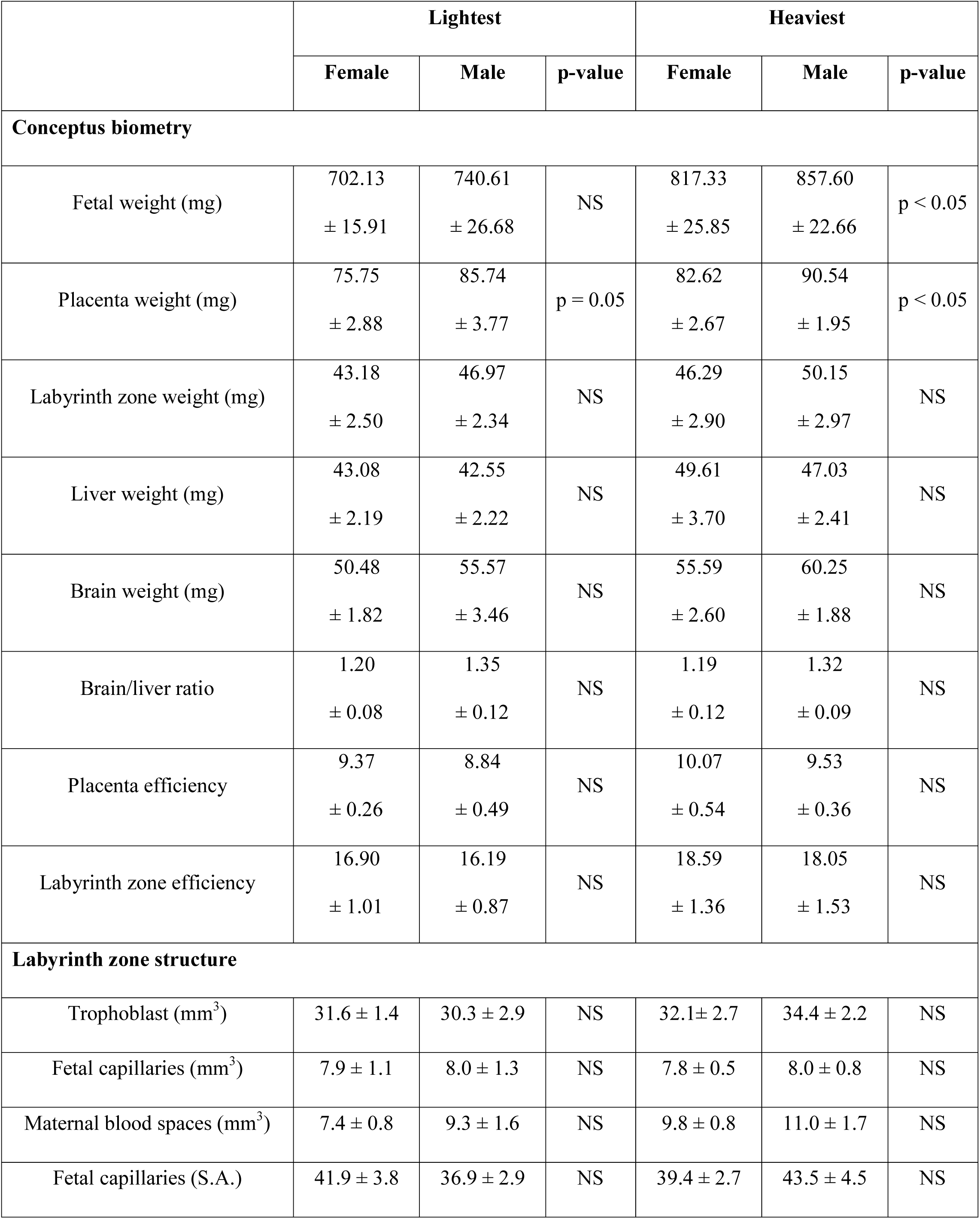
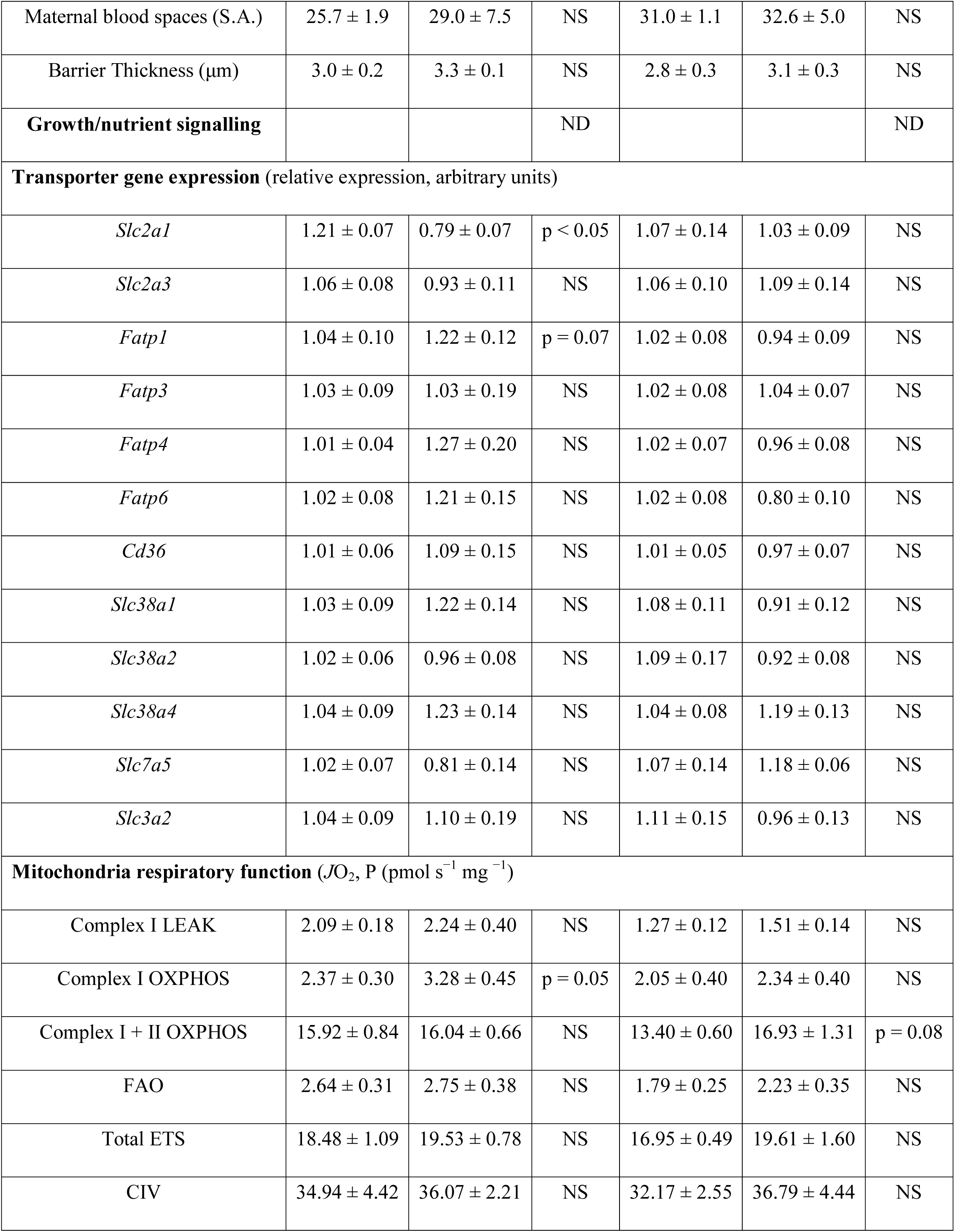
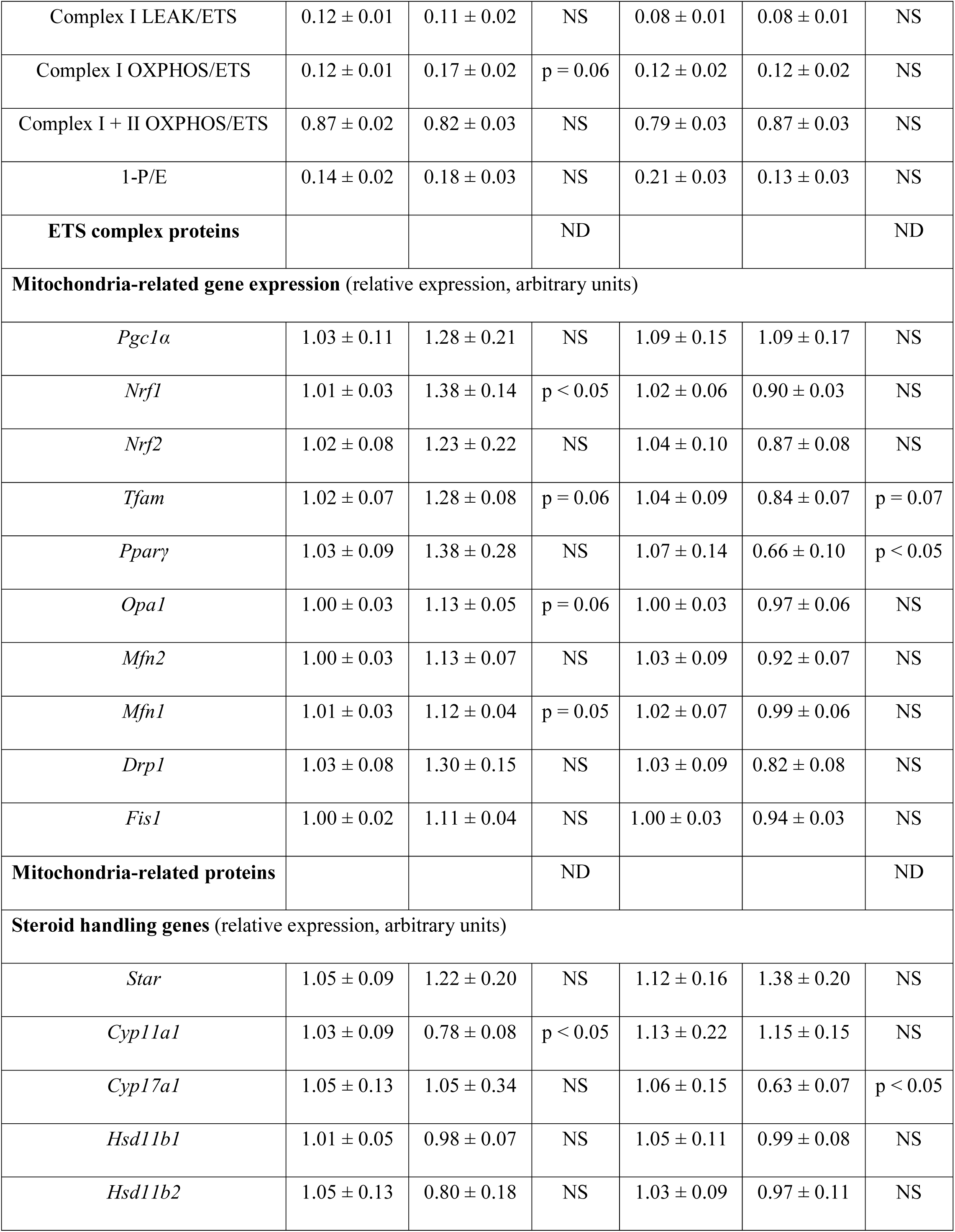
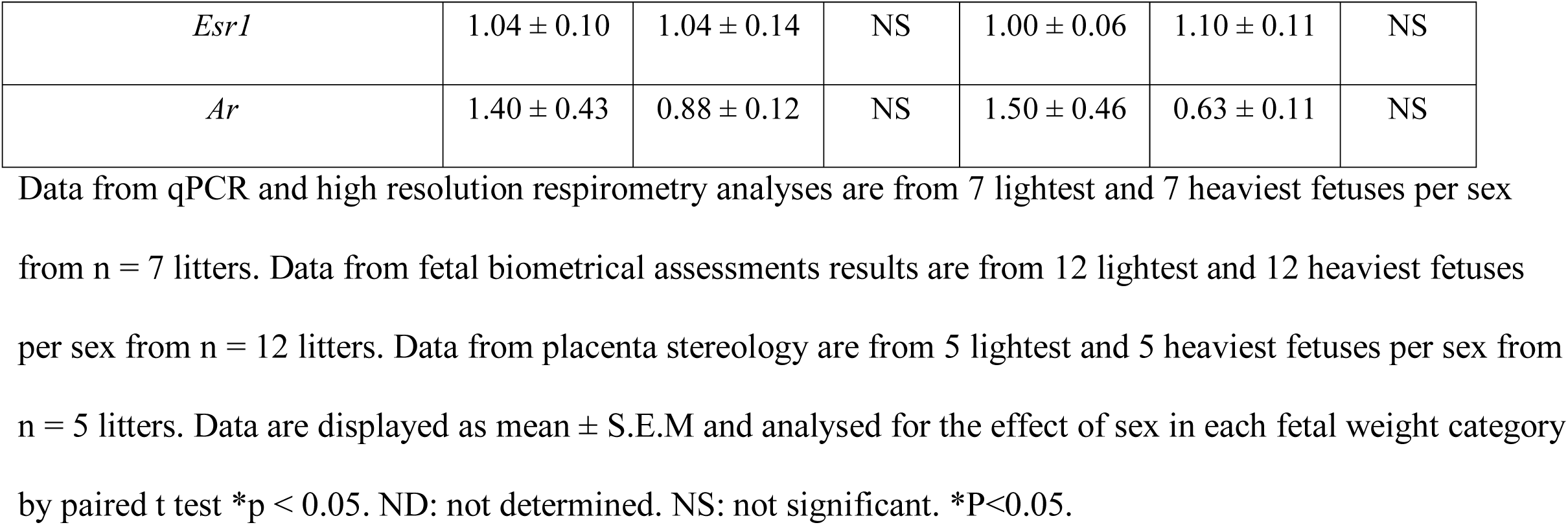
Comparisons between fetal sexes on lightest and heaviest fetuses.

## Discussion

In line with other work, this study in mice shows that feto-placental weight is on average, greater for males compared to females. Furthermore, depending on fetal sex, the relationship between fetal weight and placental weight varies within the litter. The principal aim of this study was therefore to understand how fetal weight differences in normal physiological mouse pregnancies relate to placental function in the two sexes separately. We show that placental mitochondrial functional capacity does indeed alter with respect to natural differences in the weight of females and male fetuses within the litter. The placental Lz of the lightest female and male fetuses showed altered abundance of ETS complex proteins and mitochondrial biogenesis genes when compared with their respective heaviest counter-parts, however, the specific nature of these changes depended on fetal sex. Moreover, the morphology, respiratory capacity, mitochondrial fission, and abundance of misfolded protein regulators of the placental Lz differed between the lightest and heaviest females, but not males. Furthermore, the level of nutrient (glucose) transporter genes varied between the lightest and heaviest males, but not females, whereas the ability to produce steroid hormones (informed by expression of steroidogenic enzyme genes) differed only between the lightest and heaviest females within the litter. There were also sex-dependent changes in the expression of hormone responsive genes, and growth and metabolic signalling proteins in the placental Lz between the lightest and heaviest fetuses. Despite these sex-related variations, the average weight differences between the lightest and heaviest fetuses were similar for both sexes. Together, these data suggest that in normal mouse pregnancy, placental structure, function and mitochondrial phenotype appears to respond differentially to the genetically determined growth demands of the female and the male fetus (Figure 8).

**Figure 8.**
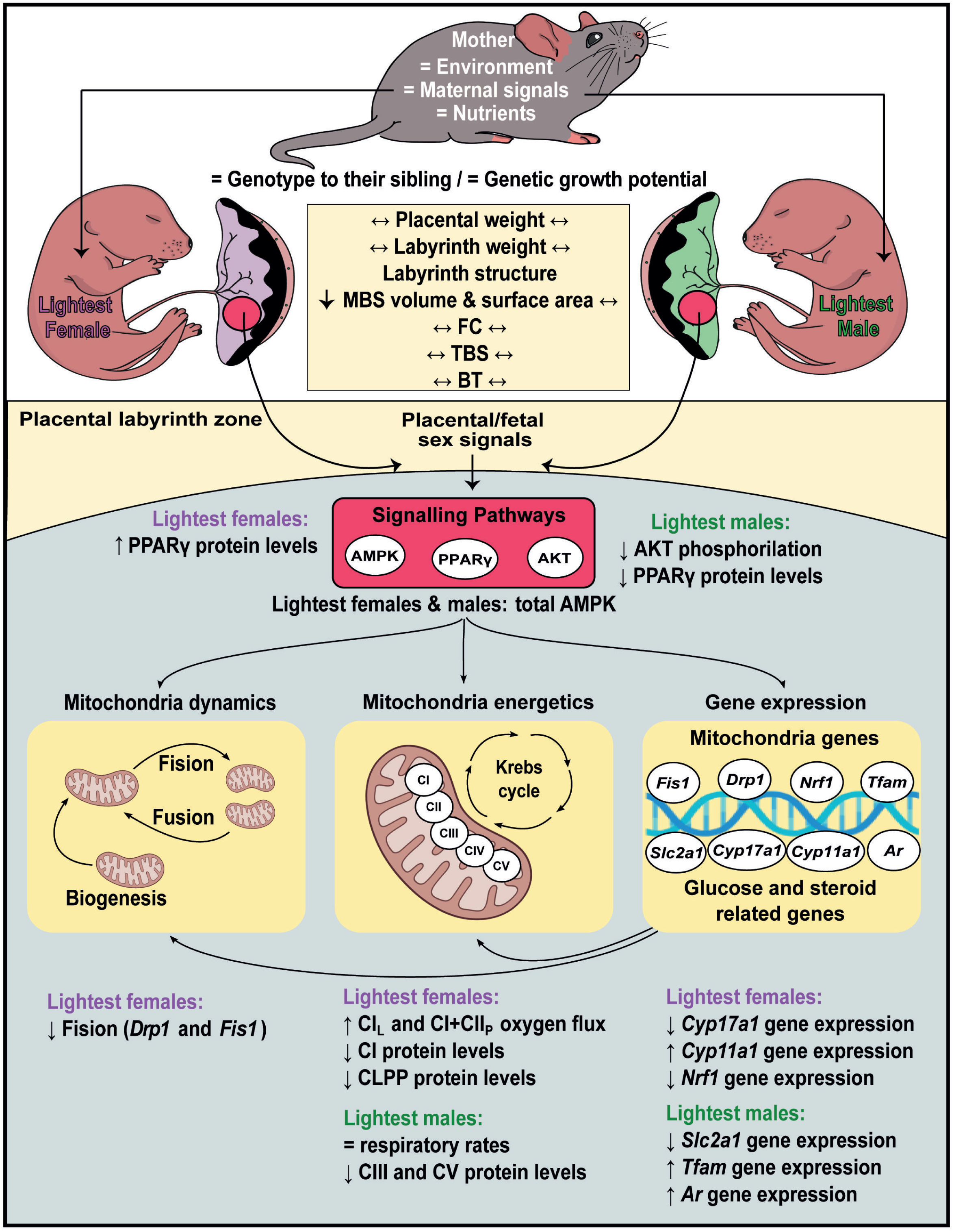
Summary figure representing the alterations in placental phenotype of the lightest *versus* heaviest fetuses of each sex in normal pregnant mice at gestational day 18. Fetuses within the litter are exposed to the same maternal endocrine and nutritional environment, yet differences in the weight of fetuses can be observed. A comprehensive analysis of the lightest and heaviest fetus from the litter revealed significant structural, functional, mitochondrial and molecular differences in the placental labyrinth zone, which in turn, depended on the sex of the fetus. We speculate that in part, these sex-dependant differences in placental phenotype between the lightest and heaviest fetuses of the litter are due to variations in the fetal endocrine and metabolic environment which operate via key signalling pathways (AMPK, AKT and PPARγ) in the placental labyrinth zone. Further work is required to provide a mechanistic explanation for sexually dimorphic differences seen and to understand their physiological relevance in suboptimal gestational environments.

## Fetal and placenta growth

In our study, placental efficiency, and other fetal biometry parameters (fractional liver and brain weights) were not different between lightest and heaviest fetuses of each sex. This is relevant since placental efficiency indicates the capacity of the placenta to support fetal growth and alterations in this measure, as well as the symmetry of fetal body growth enhances the risk for chronic diseases in later life through developmental programming [12]. Similarities in body proportionality between lightest and heaviest fetuses likely relates to the adaptative properties of their placentas in these normal, healthy pregnancies. Interestingly, previous work exploring the implications of natural intra-litter variability of placental weight, rather than fetal weight in mice, has found morphological differences between the lightest and the heaviest placentas, which included a greater Lz volume and an increased surface area for exchange [3]. Functional adaptations were also found, with a greater rate of amino acid transfer and enhanced expression of sodium-dependent neutral amino acid transporter-2 (*Slc38a2*) by the lightest *versus* the heaviest placentas [3]. Similarly, calcium transfer across lightest placenta was higher than the heaviest placentas within the litter, resulting in similar calcium accretion levels in the fetus [6]. Variations in placental structure and transport within the litter were related to an increase in placental efficiency in both of these previous studies [3, 6]. However, in our study, placentas sustaining the lightest or the heaviest fetuses were not necessarily the lightest or the heaviest placentas within the litter. In addition, a key strength of our study is that the lightest and heaviest fetuses of each sex were analysed. Segregating the data by sex identified that there was a positive correlation between placental and fetal weight for only the lightest females in the litter. These data indicate that there may be differences in the way in which the placenta may be supporting growth of the female and male fetuses within the litter. This is consistent with other work in humans suggesting that the relationship between placental weight and birth weight differs statistically between females and males and may reflect sexual dimorphism in placental reserve capacity and prioritisation of somatic growth [29].

While there was no difference in placental fetal capillaries, trophoblast volume or barrier thickness between the lightest and heaviest of either fetal sex, maternal blood space volume and placenta surface area was lower in the lightest compared to the heaviest female fetuses. These differences in placental structure suggest mal-perfusion of the placenta of the lightest females and would be expected to decrease the delivery of nutrients and oxygen to fetus, and could explain the weight discrepancy with the heaviest females in the litter. Indeed, previous studies in pigs have shown that compared to placentas supporting fetuses weighing closest to the litter mean, placentas supplying the lightest fetuses within the litter have impaired angiogenesis [30]. Moreover, work in rats has suggested that an angiogenic imbalance may underlie differences in uteroplacental vascularization and fetoplacental development within the litter [31]. It would be interesting to identify whether there are alterations in angiogenic factor expression that explain the sex-related differences in placental morphology of the lightest *versus* heaviest fetuses in the litter.

## Placental mitochondrial function

Previous work has shown that in human pregnancies associated with FGR, although mRNA expression of ETS complexes (II, III and IV) is lower, there is higher mitochondrial DNA (mtDNA) content and higher oxygen consumption related with mitochondrial bioenergetics in the placenta [32]. Similarly, FGR induced by maternal caloric restriction in rats is associated with augmented mitochondrial biogenesis, as evidenced by the increased expression of PGC-1α,, NRF1 and TFAM, as well as elevated complex I and IV dependent respiration in the placenta [20]. These data suggest that a common response of the placenta to try and meet the genetically determined demands of the fetus for growth during gestation involves the modulation of placental mitochondrial respiratory capacity. The current study supports this notion and shows that the functional characteristics of placental mitochondria also adapts with natural variations of fetal growth in normal pregnancy. In females, expression of the biogenesis promoter gene *Nrf1*, fission regulator genes *Drp1* and *Fis1*, and mitochondrial complex I protein were lower, yet complex I LEAK and complex I+II OXPHOS rates were greater in the placenta supporting the lightest compared to the heaviest females. Whereas in males, biogenesis gene *Tfam* was greater, yet mitochondrial complexes III and V proteins were lower and LEAK and OXPHOS rates were not different in the placenta supporting the lightest compared to the heaviest males. Therefore, for both sexes, the placentas of the lightest fetuses appears to increase mitochondrial respiratory efficiency (as there was reduced mitochondrial complexes yet unaltered or increased respiration), although the underlying mechanisms and extent to which this may occur is different for females and males. Indeed, our results suggest that mitochondria in the placenta sustaining the lightest female fetuses in the litter are more responsive to adaptative mechanisms, as they exhibited increased mitochondrial respiration rates. This enhanced adaptive response may have been beneficial in providing the energy to sustain the expression of glucose transporters for the lightest fetuses. Previous work has demonstrated there is reduced abundance of all mitochondrial complexes and lower OXPHOS respiration rates in placental trophoblast from obese women who deliver high birth weight babies [33]. Moreover, mitochondrial complex activity is also decreased in the placenta from women with pre-pregnancy obesity or pre-gestational diabetes who have LGA babies [34]. Thus, the natural variation in intra-litter placental mitochondrial function in the current study is likely the outcome of adaptive responses in operation for both the lightest and heaviest fetuses.

In the lightest female, but not lightest males, there was lower CLPP protein abundance when compared to the heaviest female fetuses of the litter. CLPP is also decreased along with mitochondrial complex abundance in the placenta of preeclamptic women delivering FGR babies [35]. The biological relevance of the sex-specific difference in placental CLPP level between the lightest and heaviest fetuses is currently unknown. However, differences in placental CLPP protein may be particularly relevant for the outcome of female and male fetuses if the gestation is challenged, such as by a hypoxic or nutritional stimulus [23].

## Placental sex steroid handling

The placental expression of key steroid synthetic enzyme genes was differentially altered between the lightest and heaviest females only. The greater *Cyp11a1* while lower *Cyp17a1* gene expression in the lightest compared to the heaviest female fetus, would be expected to enhance the synthesis of the steroid hormone precursor pregnenolone, but also limit the synthesis of sex steroids. CYP11A1 and CYP17A1 proteins are both cytochrome P450 monooxygenases located in the mitochondrial membrane that use oxygen for steroidogenesis, and changes in their expression may have relevance for understanding the greater rate of oxygen consumption in LEAK state for the placenta of the lightest *versus* heaviest females. Other work has shown that CYP11A1 protein is upregulated in the placenta of women with preeclampsia and overexpression of CYP11A1 protein in human trophoblast cells reduces proliferation and induces apoptosis [36, 37]. In addition, *in vitro* studies using cell lines have implicated an important role of CYP17A1 in placental estrogen production [38]. Thus, further work would value from quantifying steroid hormone levels in the placenta. The expression of the androgen receptor gene was greater in the placenta of the lightest compared to the heaviest males; a difference not seen in females. These data suggest enhanced sensitivity of the lightest male placenta to androgens, namely testosterone, which can be produced by the fetal testes from approximately day 12-13 of mouse [39]. Interestingly, in rats, elevated testosterone levels disrupts the number and structure of mitochondria in the placenta and decreases fetal weight [40, 41]. Additionally, DHT (5α-reduced metabolite of testosterone) and insulin treatment of rats induces mitochondrial damage and an imbalance between oxidative and anti-oxidative stress responses in the placenta in association with FGR [42]. Thus, differences in steroid production and signalling are likely involved in the underlying alterations in placental morphology and mitochondrial functional capacity supporting the lightest fetus of each sex.

## Placental signalling pathways

The mechanisms underlying the differences in intra-litter placental mitochondrial function for each fetal sex are unknown. However, intra-litter differences in placental morphology and mitochondrial functional capacity likely stem from variations in the abundance of AMPK, AKT and PPARγ proteins between the lightest and heaviest female and male fetuses [15]. Increased levels of AMPK protein in the placenta were seen for both the lightest females and lightest males compared to their heaviest counterparts. AMPK is activated by an increase in the AMP to ATP ratio and hence, is reflective of a decline in energy status. In turn, AMPK activates metabolic enzymes that allow cells to switch on catabolic pathways that generate ATP, including glycolysis and fatty acid β-oxidation [43]. We did not observe differences in placental fatty acid oxidation between the lightest and heaviest of each fetal sex within the litter. However, it would be beneficial to assess glycolysis and glycolytic enzyme expression in the placenta to assess whether there are intra-litter differences for females or males. In addition to its role in energy sensing, AMPK regulates placental trophoblast differentiation, proliferation and nutrient transport [44]. Moreover, AMPK in the placenta has been linked to maternal vascular responses and changes in placental morphology and fetal growth in hypoxic pregnancy [8, 45]. Interestingly, compared to their heavier counterparts, the magnitude of increase in placental AMPK protein was greater for the lightest females *versus* for the lightest males (increased by 67% and 41%, respectively). Whether this may relate to the observation of altered placental morphology in only the lightest females requires further study.

Activation of AKT (phosphorylated AKT protein) was lower in the Lz of the lightest compared to heaviest male fetuses only. The AKT-mTOR signalling pathway plays a crucial role in the regulation of placental transport function and it has been shown to be up-regulated in pregnancies from obese women delivering LGA babies [46] and down-regulated in placentas from SGA/FGR babies [47]. Moreover, placental trophoblast specific loss of phosphoinositol-3 kinase (PI3K) signalling, which is upstream of AKT, leads to FGR in mice [48]. In line with the reduced AKT activation, only the lightest males presented lower glucose transporter gene expression (*Slc2a1*, encodes GLUT1) in their placenta, when compared to the heaviest males. Insulin is a major fetal growth factor that signals via AKT to mediate its metabolic effects. Reduced activation of AKT in the placenta of the lightest males may therefore reflect reduced fetal insulin signalling to the placenta [49]. These data may also reinforce the idea that the placentas of females and males execute their own molecular mechanisms to best support fetal growth and development. In humans, placental GLUT1 protein is down-regulated in preeclampsia, and this might play a role in the coincident development of FGR [50]. Conversely, women with insulin-dependent diabetes who have an increased incidence of LGA show increased GLUT1 protein expression and a higher mediated uptake of D-glucose by the placenta [51]. Since glucose is the most important energetic substrate for fetal growth, lower expression of placental *Slc2a1* gene in the lightest relative to the heavier male fetus may explain their fetal growth discrepancy.

Sexually dimorphic differences in placental phenotype may also relate to changes in PPARγ protein abundance; PPARγ was greater in the lightest females, but lower in the lightest males compared to their respective counterparts. In humans, PPARγ expression was found to be reduced in the placenta of SGA fetuses and to associate positively with fetal and placental weights within this subpopulation [52]. In addition, it has been reported that PPARγ modulates the expression of amino acid transporters LAT1 and LAT2 (encoded by *Slc7a5* and *Slc3a2* genes, respectively) and plays a key role in the control of fetal growth [53]. PPARγ is also important for the regulation of lipid uptake and metabolism and regulates mitochondrial function via multiple routes [54]. While no differences were detected in the placental expression of amino acid and fatty acid transporters, changes in PPARγ protein could in part mediate the differences in mitochondrial respiratory capacity and regulatory factor expression between lightest and heaviest fetuses in our study. PPARγ is a transcription factor that can be modulated by numerous signals, including hormonal/growth factor signalling pathways, inflammatory/stress signalling pathways and cellular metabolite levels [55]. Hence, the sex-related alterations in PPARγ abundance in the placenta of the lightest females and the lightest males likely reflects variations in the metabolic and hormonal environment of those fetuses relative to their heaviest counterparts within the litter.

## Placental phenotype and fetal sex comparisons

Previous work has shown there are ontogenic changes in placental Lz morphology, function, mitochondrial respiration and mitochondrial-related regulators that support the growth demands of the fetus during normal late mouse pregnancy [23]. In the current study the heaviest males and their placentas were heavier than the heaviest females within the litter, although no differences were found in placental morphology, mitochondrial respiratory capacity (respiration rate or mitochondrial-related gene expression), or transport/hormone genes between them. In contrast, the lightest males and their placentas did not differ in weight when compared to the lightest females, yet they varied in the placental expression of nutrient transporters, steroidogenesis genes, mitochondrial respiration (complex I OXPHOS rate) and mitochondrial-related gene expression. These data suggest that male and female fetuses differentially execute a placental response depending on their ability to reach (lightest fetuses) or supersede (heaviest fetuses) their genetic growth potential. Assessing fetal hormone and nutrient/metabolite levels in the lightest and heaviest fetuses of the litter may provide some insight into the mechanisms underlying the sexually dimorphic differences seen in the placenta. The possible involvement of fetal hormone and nutrient/metabolites in mediating adaptations in the placenta could be tested using fetal specific manipulations [48], but identification of the precise underlying mechanisms may be highly challenging. Future work would also benefit from assessing the timing of changes occurring in the placenta relative to the pattern of fetal growth for the males and females within the litter. This will help to identify whether fetal weight discrepancies within the litter are the cause or consequence of placental adaptations that started during early mouse pregnancy.

## Study strengths and limitations

Our study has clear strengths. It provides a comprehensive analysis of the structural, functional, mitochondrial and molecular differences in the isolated transport labyrinth zone for the lightest and heaviest fetuses of each sex in litters of normal, healthy pregnant mice. However, the labyrinth zone is composed of numerous cell types, and the contribution of each cell population to the specific placental alterations seen is unknown. We also do not know if there are alterations in the endocrine junctional zone, which is also important for the support of fetal growth and can exhibit sexual dimorphism [56]. Moreover, as did not record uterine position of the individual fetuses, whether placental changes are driving alterations in fetal growth based on maternal supply differences secondary to perfusion/implantation variations in each sex could not be ascertained. Indeed, differences in the placenta could also be influenced by sex of adjacent fetuses and variations in litter size [31,57,58], which would need to be addressed using much larger sample sizes to ensure there is sufficient statistical power.

## Summary

In summary, our data show that the placental transport zone (Lz) adopts different strategies, at the level of morphology, nutrient transport, steroid handling, and mitochondrial function to support growth of the lightest and the heaviest fetuses within the litter in normal physiological mouse pregnancy. These adaptations are likely mediated via metabolic (e.g. lipids, energy status) and endocrine cues (insulin, sex steroids) within the fetus that trigger signalling pathways (e.g. AMPK, PPARγ, AKT) in the placenta, initiating pleiotropic effects. Further work is required to test the mechanisms underlying phenotypic differences in the placenta and to ascertain the relevance of our findings for pregnancies with adverse conditions, such as maternal malnutrition, obesity, or reduced oxygen availability where the maternal ability to provide resources to the fetus for growth are constrained. From a clinical perspective, our data may be important for understanding the pathways leading to placental insufficiency and fetuses not reaching (FGR/SGA) or exceeding their genetically determined growth potential (LGA). They may also have significance in understanding the discordance in weight and perinatal outcomes between babies of multiple gestations in women. Moreover, since the spectrum of pregnancy outcomes and the factors causally involved are likely to be many, determining how placental phenotype interacts with the weight of female or male fetuses within normal mouse litter may be useful to the design of sex-specific therapeutic agents to improve pregnancy outcomes in humans. This is highly relevant given the profound impacts of fetal growth and pregnancy complications on immediate and life-long health of the child.

## Funding

ESP was supported by a Beca-Chile, ANID Postdoctoral Scholarship: 74190055. DPC was supported by Programa Institucional de Internacionalização (PRINT) [Grant number 88887.508140/2020-00]. JLT currently holds a Sir Henry Wellcome Postdoctoral Fellowship [Grant number 220456/Z/20/Z] and previously a Newton International Fellowship from the Royal Society [NF170988 / RG90199]. ANSP is supported by a Medical Research Council New Investigator grant and Lister Institute of Preventative Medicine Research Prize (MR/R022690/1 / RG93186 and RG93692, respectively).

## Author’s contribution

ESP, JLT and ANSP designed the study. ESP, JLT and DPC performed the experiments and analyzed and graphed the data. ESP and ANSP wrote the paper. All authors contributed to data interpretation and performed final editing checks and approved the final manuscript.

## Conflicts of interest/Competing interests

The authors declare that no conflicts of interest/competing interests exist.

## Data availability

All data are available upon reasonable request.

---

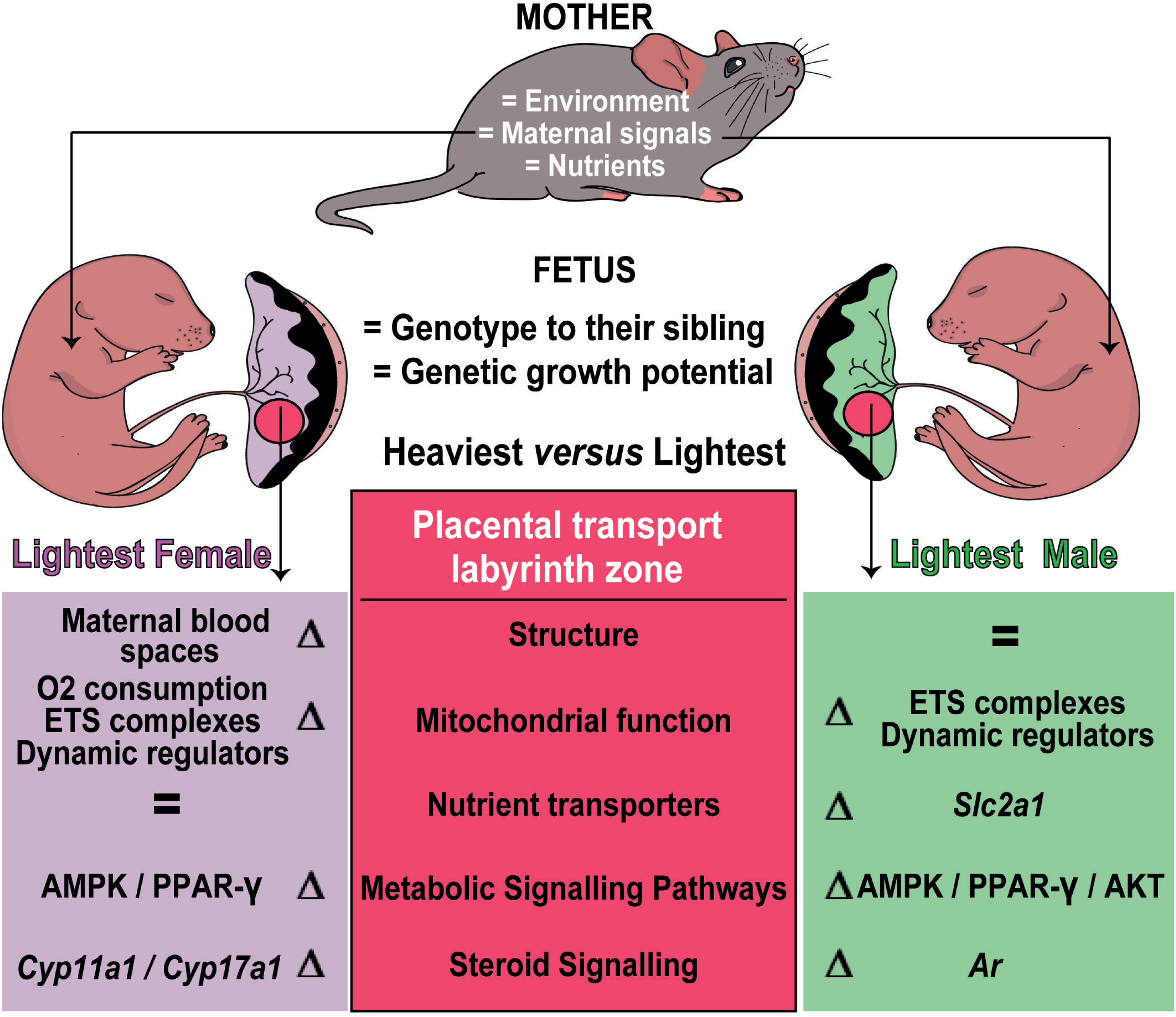

## Notes

### Competing Interest Statement

The authors have declared no competing interest.

